# Redefining hematopoietic progenitor cells and reforming the hierarchy of hematopoiesis

**DOI:** 10.1101/2023.01.27.524347

**Authors:** Lipeng Chen, Qing Sun, Guoqiang Li, Qijun Huang, Sujin Chen, Yingyun Fu, Yongjian Yue

## Abstract

Deciphering the mechanisms underlying progenitor cell differentiation and cell-fate decisions is critical for answering fundamental questions regarding hematopoietic lineage commitment. Here, we redefine the entire spectrum of original hematopoietic progenitor cells (HPCs) using a comprehensive transcriptional atlas that effectively delineates the transitional progenitors. This is the first study to fully distinguish the transitional state along hematopoietic progenitor cell differentiation, reconciling previous controversial definitions of common myeloid progenitors (CMPs), granulocyte–monocyte progenitors (GMPs), and lymphoid-primed multipotent progenitors (LMPPs). Moreover, plasma progenitor cells are identified and defined. Transcription factors associated with key cell-fate decisions are identified at each level of the hematopoietic hierarchy, providing novel insights into the underlying molecular mechanisms. The hematopoietic hierarchy roadmap was reformed that reconciles previous models concerning pathways and branches of hematopoiesis commitment. Initial hematopoietic progenitors are simultaneously primed into megakaryocytic–erythroid, lymphoid, and neutrophilic progenitors during the first differentiation stage of hematopoiesis. During initial progenitor commitment, *GATA2*, *HOPX*, and *CSF3R* determine the co-segregation of the three transitional lineage branches. Two types of lineage-commitment processes occur during hematopoiesis: the megakaryocytic–erythroid lineage commitment process is continuous, while the lymphoid-lineage commitment is stepwise. Collectively, these results raise numerous possibilities for precisely controlling progenitor cell differentiation, facilitating advancements in regenerative medicine and disease treatment.

**Highlights:**

- Hematopoietic progenitors are redefined using a comprehensive transcriptional atlas.
- Cell fate decision-related transcription factors are revealed in the hematopoietic hierarchy.
- Progenitor lineage commitment includes continuous and stepwise processes.
- The initial hematopoietic hierarchy is simultaneously primed into three lineages.

## Introduction

Hematopoietic stem cells (HSCs) comprise a rare cell population in the peripheral blood, capable of self-renewal and differentiation into full blood cell lineages with tree-like differentiation patterns in classical hematopoietic hierarchical models [1]. While the diversity of human peripheral blood progenitor cells is well-known, their conflicting definitions, lineage heterogeneity, and dynamic differentiation impede the characterization of the molecular mechanisms underlying lineage commitment. Moreover, the definition of hematopoietic progenitor cells (HPCs) has not been standardized due to the limitations in sorting methods and traditional definitions based on surface markers, including CD34, CD38, FLT3, and KIT [2]. Due to progenitors’ rare and transient nature, surface markers fail to efficiently capture their entire spectrum and internal state or reflect the dynamic differentiation mechanisms underlying hematopoiesis [3, 4]. Moreover, using original quiescent progenitors derived from bone marrow, which are widely used for functional studies and subgroup identification, may cause bias as they do not cover the entire spectrum of progenitor stages [5, 6]. HPCs have been characterized in functional studies using selected CD34^+^ cells; however, diverse cellular experimental conditions can lead to the inaccurate interpretation of HPC fate decisions as the conditions do not apply to all branches of lineage commitment [7]. Therefore, determining cell lineage definitions and fate decisions, as well as establishing reliable lineage commitment processes during hematopoiesis, remains challenging [8, 9].

Most established hierarchies of hematopoiesis are based on *in vitro* and *in vivo* functional experimental results [7, 10]. Single-cell RNA (scRNA) sequencing technology has been successfully implemented to explore the molecular heterogeneity and hierarchy of HPCs [11, 12]. However, the scRNA transcription atlas tends to modify aspects of the classical hematopoietic hierarchy, in which progenitors continuously differentiate during hematopoiesis [4, 13]. Lineage commitment is affected by transcription factors (TFs) within a continuum of cell fate-decision processes [2, 14]. However, CD34^+^ progenitor clusters exhibit a scattered and mixed distribution based on typical markers, which contradicts the unsupervised dimensionality reduction and clustering using the transcription atlas [12]. Additionally, the unsupervised dimensionality reduction and clustering of lineage commitment can result in a farraginous hierarchy model [12]. Furthermore, the inefficient and inaccurate classical definitions of hematopoietic progenitors can result in disorganized subgrouping without reliable boundaries [11, 15]. Indeed, several previously proposed hematopoietic hierarchical models contradict each other [8, 16]. Meanwhile, the critical functional roles of TFs in lineage commitment and the regulatory mechanisms underlying the hierarchy of hematopoiesis remain unclear [17, 18].

In this study, we aimed to redefine adult HPCs, identify key TFs involved in lineage-fate decisions, and reform the hematopoietic hierarchy. Our HPC atlas redefines the cellular properties of progenitors at different stages throughout hematopoietic cell-lineage commitment. Thus, our results provide new perspectives for characterizing, defining, and classifying HPCs. This work represents a significant step in clarifying the initial primed transitional progenitor and reforming the hematopoietic hierarchy, resolving complexities in hematopoiesis commitment pathways and models.

## Results

### Determining the characteristics of progenitor cells in adult peripheral blood via scRNA sequencing

To enrich the entire HPC spectrum and intermediate-state progenitors, mature lymphocytes and granulocytes were removed using a surface marker panel (Lineage^−^) via negative deletion (Figure 1A). The mean percentage of CD34^+^ cells reached 10.3% and 46.1% in control peripheral blood mononuclear cells (PBMC) and granulocyte colony-stimulating factor (G-CSF) mobilized peripheral blood groups, respectively (Figure 1B). We performed 10× Genomics scRNA sequencing to characterize the progenitor cells by unsupervised dimensionality reduction, and clustering analysis using the Seurat toolkit and uniform manifold approximation and projection (UMAP) analysis. Overall, 16,2579 qualified cells were captured and sequenced after filtering the doublets and low-quality cells, identifying 29 clusters (Figure 1C). Notably, the single-cell expression atlas showed that several CD34^+^ progenitor clusters are gathered together and present heterogeneity.

**Figure 1.**
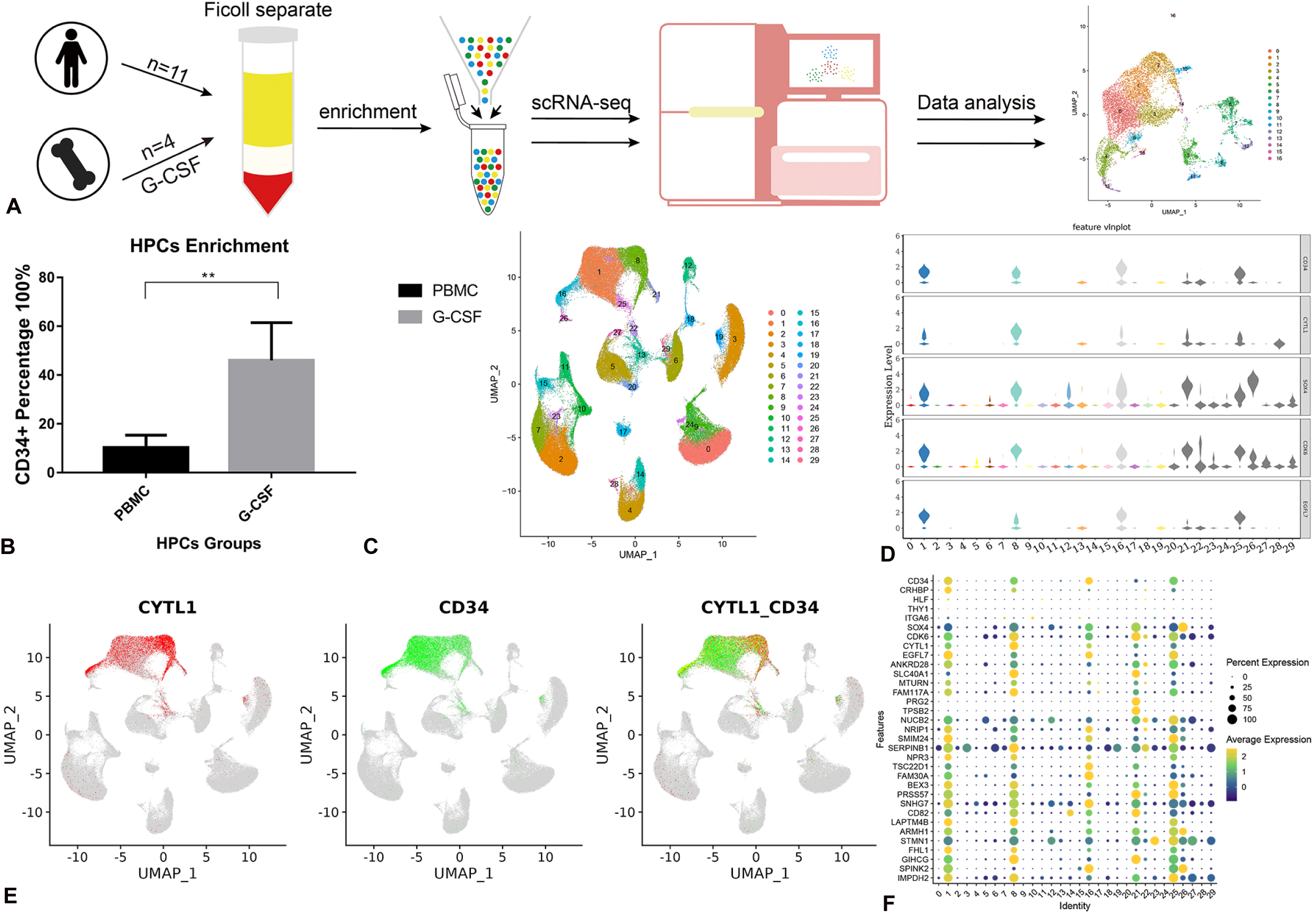
Progenitor subtypes identified in adult peripheral blood using unsupervised dimensionality reduction and clustering analysis. A, Schematic showing the progenitor cell enrichment and sequencing workflow. B, Mean percentage of CD34-positive cells in control (PBMC) and G-CSF mobilized peripheral blood groups. C, Twenty-nine integrated clusters were identified via UMAP analysis. D, Violin plot of the identified progenitor cell marker genes *CDK6*, *EGFL7*, *CYTL1*, and *SOX4*. E, Expression-feature plots of the progenitor markers *CD34* and *CYTL1*. F, Dot plot illustrating the expression of progenitor cell-related marker genes in all clusters.

Furthermore, we analyzed significantly differential genes identified among progenitor cells and other cell clusters. *CD34, SXO4*, *CDK6*, *CYTL1*, and *EGFL7* were highly expressed across the progenitor clusters (Figure 1D, E). However, the classical markers of progenitors, namely *HLF*, *THY1* (CD90), and *ITGA6* (CD49f), were negatively expressed in most progenitor clusters, excluding CRHBP (Figure 1F and S1A, B). *EGFL7* and *CYTL1* were highly and specifically expressed in the progenitor clusters accompanying expression differences among lineages, indicating that they likely represent unique marker genes for the HPC lineage. The 30 most highly expressed genes in progenitor cells are presented in Figure 1F and may have essential roles in maintaining progenitor functions and stemness. Six highly specific HPC genes were identified, namely *EGFL7, CYTL1, LAPTM4B, PRSS57, NPR3*, and *SMIM24*. Combining these marker genes achieved high efficiency in identifying and characterizing HPCs. Gene Ontology (GO) and pathway analysis further revealed the enrichment of immune activity and regulation of hematopoietic cell lineages in HPC clusters (Figure S1C, D). Gene set enrichment analysis of differentially expressed genes showed that the hematopoietic cell lineage, MAPK, and PI3K–Akt pathways were enriched in progenitor cells, but not significantly (Figure S1E). Therefore, based on the identified conventional and novel markers and the aforementioned key functional genes associated with HPCs, we successfully characterized the identified CD34^+^ HPC clusters.

### Conventional surface markers are ineffective at accurately defining HPCs

To reveal the heterogeneity of HPCs, we first extracted and re-clustered the aggregated HPC subsets. Results showed that *CYTL1* and *CD34* negative cells were clustered in the same subgroup, indicating that they were not progenitor cells (Figure S2A). Notably, the expression characterization of *EGFL7, CYTL1, LAPTM4B*, and *NPR3* was highly aggregated in CD34 positive clusters and presented as more efficient and sensitive than CRHBP or HLF in progenitor cell identification (Figure S2B). Given that the expression of CD34 was more uniform and pronounced than that of other markers, further unsupervised dimensionality reduction re-clustering analysis was performed on 25,450 CD34^+^ cells to evaluate diversity, resulting in 14 clusters, including 1,9953 mobilized and 5,497 quiescent progenitors (Figure 2A). A sufficient number of progenitor cells were present in each cluster to generate a reliable transcriptional atlas (Table S1). These progenitor cell subgroups exhibited categorical and functional heterogeneity, clustering in three main differentiation directions (Figure 2A). Conventionally, membrane markers of common myeloid progenitors (CMPs) and granulocyte–monocyte progenitors (GMPs) are defined based on the CD38^+^MME^−^FLT3^+^CD45RA phenotype, whereas megakaryocytic–erythroid progenitors (MEPs) are defined according to the CD38^+^MME^−^FLT3^−^ phenotype [2]. However, our results showed that CD38, FLT3, IL3RA, and KIT are pan-expressed in most progenitor clusters and were indistinguishable among the subgroups (Figure 2B, C). The FLT3^+^ progenitor cells were a diverse population, which may explain the varying differentiation potentials observed in previous functional studies of select FLT3^+^ progenitors [19, 20]. Moreover, *GATA1* and *FLT3* showed opposing expression patterns in the clusters (Figure 2B), indicating that CD34^+^GATA1^+^ and CD34^+^FLT3^+^ progenitors represented two distinct differentiation directions. Canonical markers, including *CD49f* (*ITGA6*) and *THY1*, were also expressed at very low levels in HPCs (Figure 2C). However, the TFs *GATA1* and *GATA2* showed high effectiveness in distinguishing HPC intermediate stages (Figure 2B).

**Figure 2.**
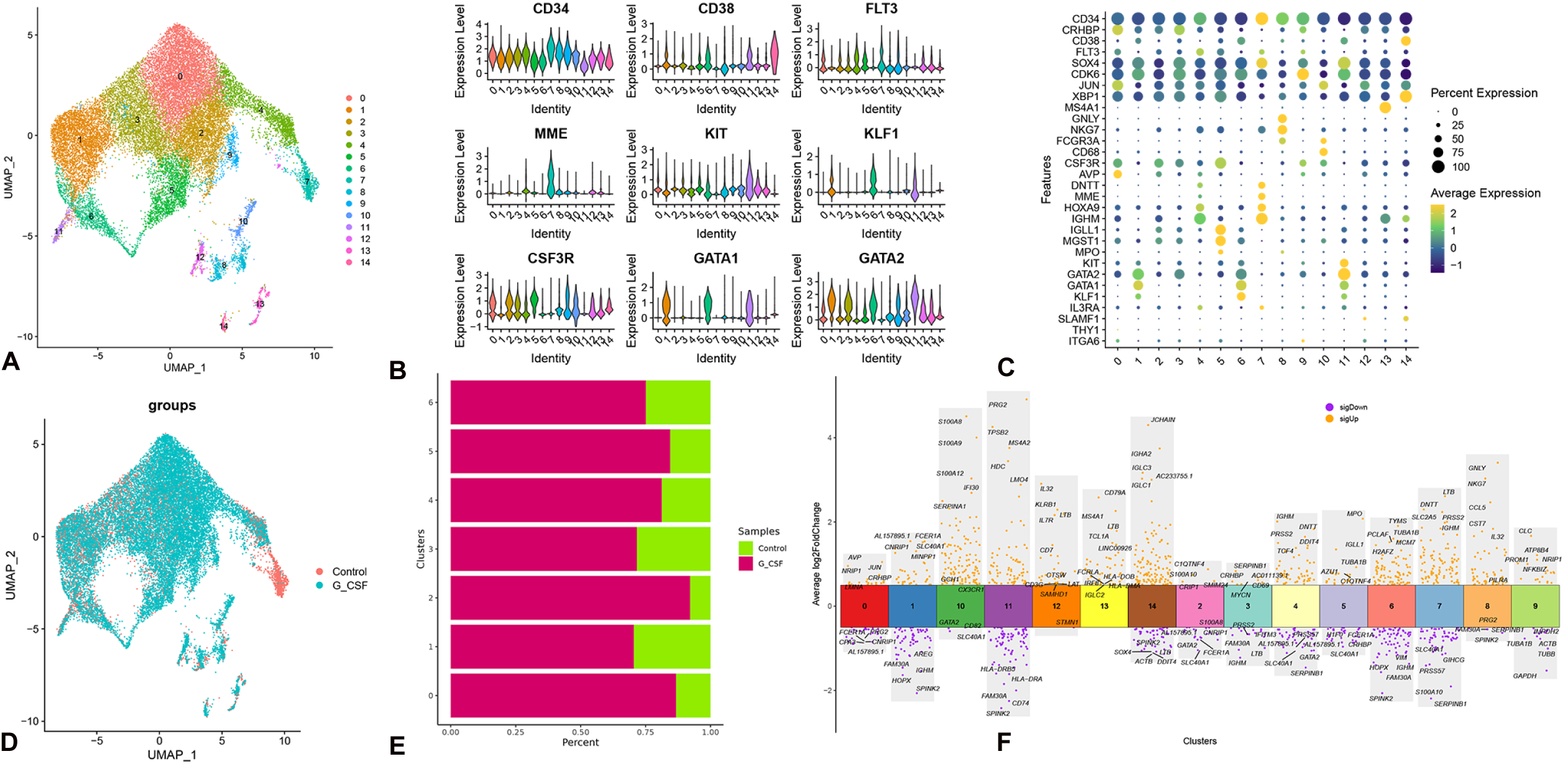
Characterization of progenitor subtypes in unperturbed and activated states. A, UMAP illustrating 14 clusters of integrated CD34^+^ progenitor cells. B, Violin plots showing the expression of conventional surface markers and unique marker genes, indicating that *GATA2*, *CSF3R*, and *MME* were more efficient and accurate markers than the conventional surface markers, *KIT* and *FLT3*, for classifying myeloid, granulocyte, and lymphoid progenitors. C, Dot plots showing the expression of representative markers and lineage-specific genes. D, UMAP showing the distribution of cells in the unperturbed (Control) and activated (G-CSF) states, displaying the persistence of all types of progenitors in the quiescent state of adult peripheral blood. E, Bar plot displaying the cell number and proportion changes in all progenitor clusters. The primitive clusters (C0) exhibited the most pronounced changes in terms of cell numbers and fold changes. F, Volcano plot showing the top five genes with the highest fold-changes in expression among the progenitor clusters. C0, cluster 0; UMAP, uniform manifold approximation and projection.

We subsequently merged a bone marrow single-cell cohort of 7,037 CD34^+^ cells from seven projects to validate the identified markers. The bone marrow progenitor cells exhibited scattered expression distributions, potentially caused by the inefficient capture of the intermediate state, and an insufficient number of cells in each cluster (Figure S2C). Expression of the canonical markers *THY1, CD49f*, and *HLF* also exhibited low efficiency. Similarly, clusters in bone marrow CD34^+^ cells could not be clearly distinguished according to the traditional classification criteria of *FLT3, IL3RA*, and *KIT*. In contrast, the newly identified markers *EGFL7*, *CYTL1*, and *SOX4* displayed high efficiency in classifying clusters (Figure S2D), supporting the reliability of the marker identification (Figure 1E, F). Overall, our data show that using the classical and canonical markers *CD49f*, *THY1*, *FLT3*, *IL3RA*, and *KIT* to classify or define progenitor cell groups is relatively arbitrary and subjective. This leads to a heterogeneous mixture of various progenitors that are ineffectively distinguished as CMPs, GMPs, and CLPs [18, 21].

### Redefinition of adult peripheral blood progenitor cells

To accurately and efficiently redefine the progenitor clusters, new marker genes, especially TFs for distinguishing progenitor clusters, were identified in the reliable transcriptional atlas. The subset distribution reproducibility across samples was exemplary, indicating the reliable repeatability of each atlas (Figure S3A). The clustered progenitors were redefined by identifying the unique markers of each subset (Figure 2C). Expression of the canonical markers *CD49f* (*ITGA6*), *SLAMF1,* and *THY1* was absent or very low in all HPC clusters. *AVP* and *CRHBP* were expressed in HSCs and, along with *CSF3R*, were primarily expressed in cluster 0 (C0). Hence, the cluster likely comprised the original HSC progenitors designated multipotent progenitor cells (MPCs; Figure 2B and C). Myeloid progenitors were detected in C1, C3, C6, C2, and C5 with gradual upregulation of *CD38, GATA2*, *NFE2*, and *KIT* (Figure 2B). The unidirectional lineages C1, C3, and C6 could be distinguished based on the expression levels of TFs. For example, expression of the erythrocyte fate decision TFs *KLF1*, *GATA1*, and *GATA2* (Figure 2B) indicated the presence of megakaryocytic–erythroid (MkE) lineage progenitors. In contrast, C2 and C5 expressed myeloperoxidase (*MPO*), *IGLL1*, and *CSF3R* marker genes (Figure 2C). MPO is the most abundant and uniquely expressed protein in human neutrophils, with a vital role in neutrophil antimicrobial responses [22, 23]. Hence, the progenitor branch was likely neutrophil–monocyte progenitors (NMPs), with an entirely unique orientation from megakaryocyte-erythrocyte and lymphoid progenitors. In contrast, *IGHM*, *MME*, and *HOXA9* [24] were expressed in C4 and C7, confirming that this cluster comprised common lymphoid progenitor cells (Figure 2C). Expression of the TF *HOXA9* increased sequentially from C4 to C7. The characterization of CSF3R and GATA2 expression clearly demonstrated the co-segregation among NMPs, megakaryocyte-erythrocyte, and lymphoid progenitors (Figure 2B). Accordingly, TF expression fully distinguished the intermediate state and reflected the mechanisms underlying hematopoietic lineage commitment. Assessing the expression and activation of TFs with unique marker genes was efficient for defining HPC lineages.

To assess differences between mobilized and unperturbed progenitor cells, cell-distribution analysis was performed. The total number of progenitor cells was remarkably increased after G-CSF mobilization. A comparison of the progenitor cells before and after mobilization revealed that all progenitor cell subtypes existed in the unperturbed state (Figure 2D). *CSF3R* expression is consistent with G-CSF mobilization in the peripheral blood during HSC transplantation [25]. This signifies that the entire spectrum of progenitor cells exists in an unperturbed state within the peripheral blood, consistent with the cluster distribution in the mobilized state (Figure 2D). This might also indicate that unperturbed adult hematopoiesis depends on replenishment from multiple progenitor cell types. As the progenitor cells gradually differentiate into mature cells in the peripheral blood, we posited that the number and proportion of more primitive progenitors would be higher than those of mature progenitors. The largest relative abundance and increase in absolute number were found in C0 (Figure 2E), the original multipotent cluster, which can differentiate into multiple lineages.

To identify the CMPs, all early priming-stage clusters were re-clustered. A common intermediate progenitor stage was not detected among the priming-stage progenitors, and the clusters could not be distinguished based on FLT3, CD38, and IL3RA expression (Figure S3B). Hence, defining the CMPs or GMPs by surface markers would create a heterogeneous mixture of various progenitors (Figure S3C). The original MPCs were simultaneously primed to three transitional lineages, which committed to lymphoid, neutrophil/monocyte, and megakaryocytic–erythroid progenitor cell types; they subsequently matured into different hematopoietic lineages (Figure 2A). The subsets of the three transitional lineages were adjacent, lacking clear boundaries, indicating the continuous differentiation trajectories of the early commitment process. Moreover, the three transitional lineages were co-segregated simultaneously, contradicting the conclusion that lymphoid-primed multipotent progenitors (LMPPs) are co-segregated between neutrophils/monocytes and lymphoid cells [4, 15]. Although the surface markers (CD38-CD10+) of CLPs are consistent with LMPPs, they should not be defined as lymphoid-primed multipotent progenitors as they only present lymphoid-primed potential, not neutrophil/monocyte-primed potential. Rather, LMPPs (CD38-CD10+CSF3R+) likely represent a heterogeneous mixture of neutrophil and lymphoid transition progenitors [15]. Based on the successful capture of the entire spectrum of adult peripheral blood progenitor cells, including those in mobilized and quiescent progenitor states, we generated a comprehensive transcriptional atlas for adult peripheral blood progenitor cells.

To accurately redefine the HPC subgroups, we analyzed the characteristics of various progenitor cell subtypes, identifying unique and specific marker genes and key TFs of progenitor cells in each lineage (Table 1 and Figure 2C, F). Additionally, negatively expressed genes were identified to distinguish each similar or identically redefined subgroup (Table 1). The transcriptional atlases of the two lineages of C2 and C5 (NMPs) or C4 and C7 (CLPs) exhibited remarkable comparability, with C2 and C4 presenting an early tendency toward committed NMPs or lymphoid progenitor cells. We redefined the three MPC-primed intermediate directions of CLPs, *GATA* gene-controlled progenitors (GAPs), and NMPs that could not be characterized based on CD38, FLT3, or KIT expression (Figure 4A and Table 1). Some of the specific marker genes for each progenitor lineage overlapped with canonical marker genes of classically defined hematopoietic cells, providing complementary evidence of the reliable redefinition (Figure 2F).

**Table 1.**
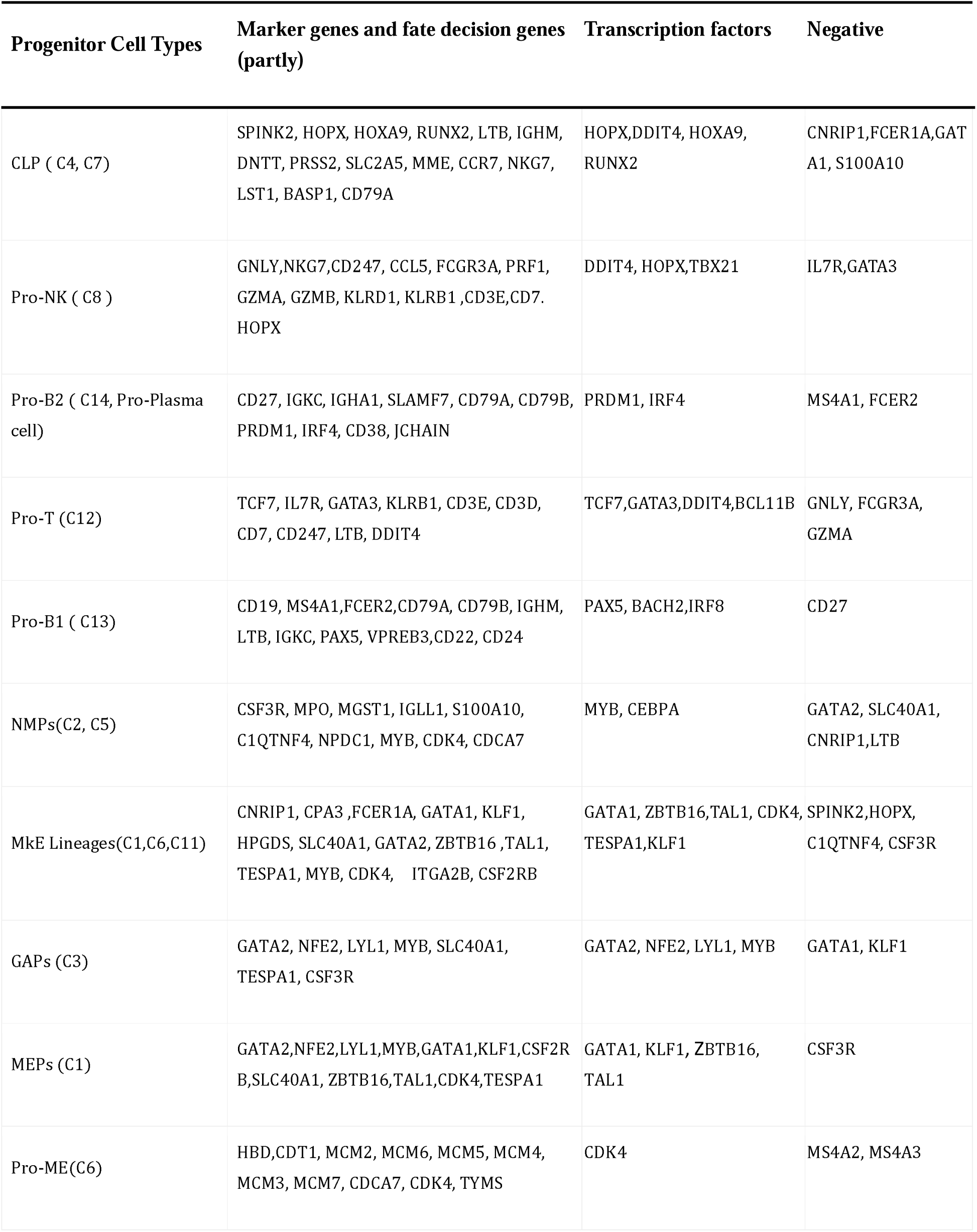
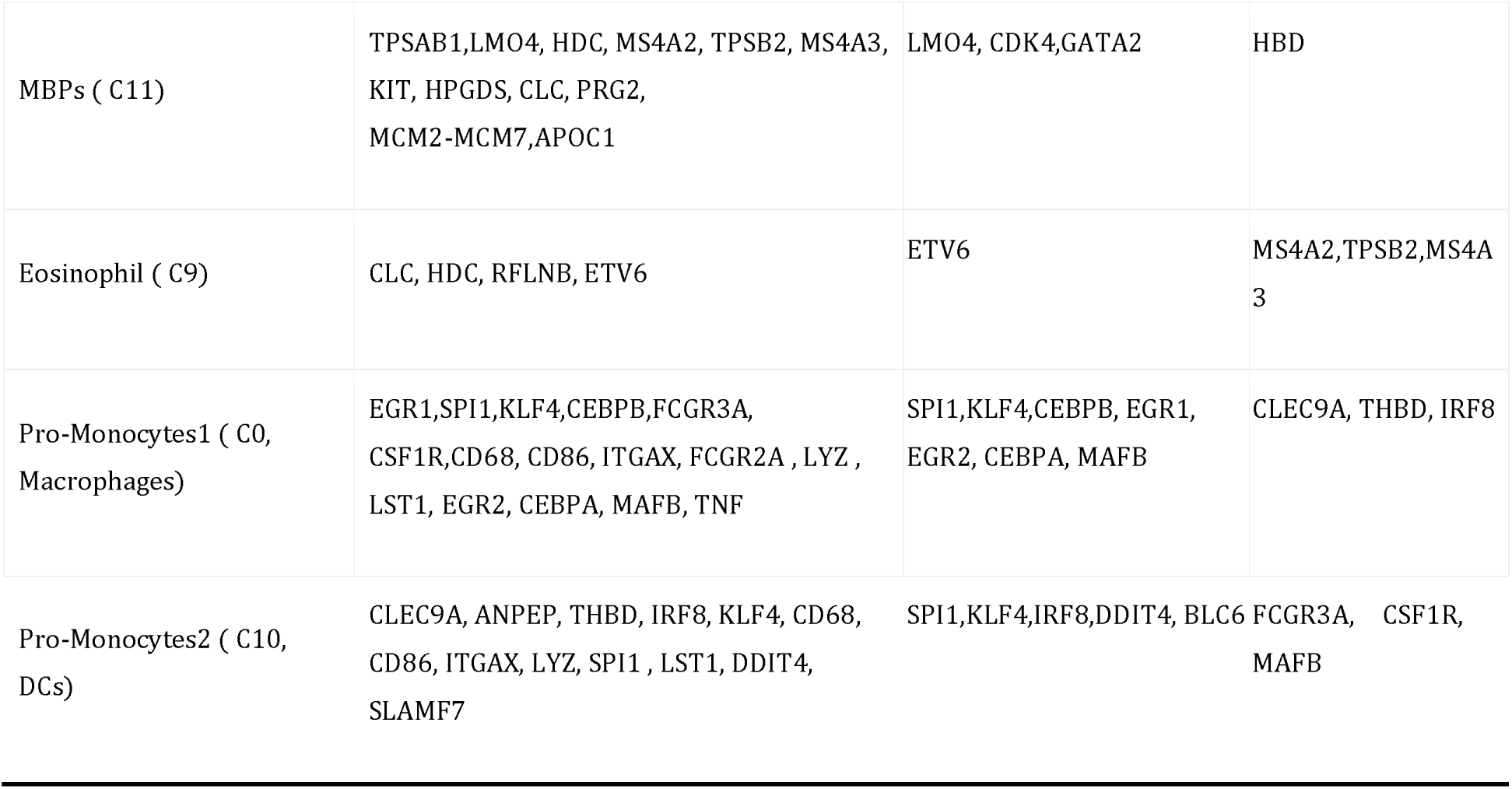
Specific marker genes and genes associated with cell-fate decisions in all redefined subgroups.

### Trajectories and dynamics of progenitor cell differentiation

As commitment trajectories can be readily revealed based on the aforementioned T-SNE distribution and reliable definition, MPCs were found to be gradually primed into three transitional progenitor cell branches and subsequently transitioned into diverse lineages (Figure 3A). To confirm the differentiation trajectories of the progenitor cells, complementary pseudo-time analysis was performed using Monocle2, which showed a consistent tendency of differentiation trajectory commitment, although the NMP lineages were a farraginous prediction (Figure S4A). The pseudo-time results showed that the redefined clusters of CLPs and MkEs were differentiated along the distinct pseudo-time trajectory states, respectively (Figure 3B, C). The dynamic changes in *GATA2* and *HOPX* expression were co-segregated and separated from each other along the lineage differentiation trajectory (Figure 3D and S4B, C). Additionally, lineage signature genes, such as *KLF1*, *GATA1*, and *GATA2* of MkEs and *HOXA9*, *HOPX*, *MME*, and *RUNX2* of CLPs, exhibited unique expression patterns along the differentiation trajectory (Figure 3E and S4E, F). The results of in-silico prediction provided complementary evidence of the progenitor differentiation trajectory. The expression trajectories of the signature genes effectively validated the reliability of the differentiation trajectories of lineages (Figure 3A).

**Figure 3.**
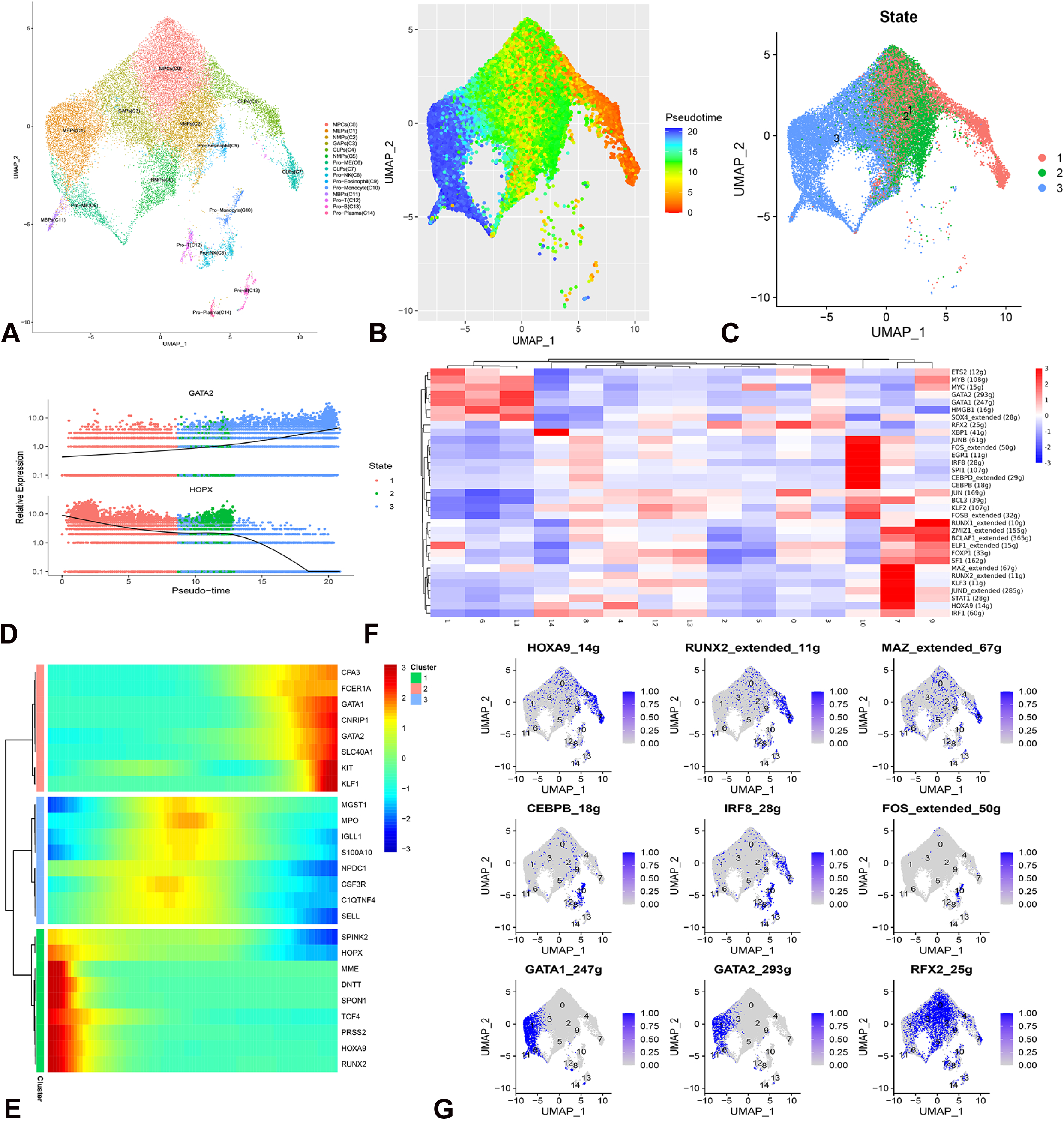
Trajectories and dynamics of progenitor cell differentiation and the activities of lineage-priming transcription factor regulons. **A,** UMAP plots demonstrating the redefined progenitor clusters, with commitment trajectories directly characterized without requiring in-silico prediction; red arrows: three simultaneously primed lineages. B, UMAP visualizing the pseudo-time trajectory scores of progenitor cells, which indicates three developmental paths from MPCs. C, UMAP feature plots of the states of pseudo-time trajectories analysis, showing that the CLP lineages were in state one (red), whereas the MkE lineages were in state three (blue). D, Plots showing the relative expression of GATA2 and HOPX across the pseudo-time trajectory. E, Heatmap showing the three distinct lineage marker gene expression patterns along the pseudo-time trajectory. F, Heatmap showing the differential activation of transcription factor regulons for all progenitor subgroups. G, Transcriptional activity of lineage signature genes related to different differentiation activation. UMAP, uniform manifold approximation and projection.

Similarly, the inference of the single-cell regulatory network and clustering assessment based on TF and regulon activities showed the activation of key TFs in cell-fate decisions during progenitor commitment (Figure 3F). The original progenitor of MPCs exhibited RFX2 activity, whereas the TFs *MAZ*, *HOXA9*, *KLF3*, *STAT1*, *JUND*, *RUNX2*, and *IRF1* were identified as key regulators of CLP priming (Figure 3G). Several TFs, including *SF1*, *RUNX1*, *ZMIZ1*, and *BCLAF1*, were identified in eosinophilic progenitors. Moreover, *GATA1*, *MYC*, *SOX4*, and *HMGB1* played critical roles in the progression of MkE lineages (Figure 3F). XBP1 was activated in plasma progenitor cells, and the TFs *IRF8*, *CEBPD*, *JUNB/FOS*, *SPI1*, *EGR1*, and *CEBPB* determined cell-fate decisions during monocytic lineage differentiation. TF and regulon activation showed a high overlap with the unique expression pattern in clusters that support cell-fate decisions in progenitor commitment. The characteristics of TF activation were similar in the same origin lineages of MkE lineages (Figure 3F). The two adjacent progenitor CLPs and the eosinophil clusters had similar TF-activation characteristics, indicating that they may have been primed by similar origin cells (C4) but still required confirmation. Therefore, integrating multi-omics and epigenetic analyses using the scRNA assay for transposase-accessible chromatin may improve our understanding of the HPC-fate-decision process. Our findings revealed highly consistent TF expression and activation patterns throughout the lineage-committing process, reflecting the synergistic effects of the suppression and activation of key regulators.

### Reforming the hematopoietic hierarchy

Based on the reliable redefinition, precise characteristics of the entire progenitor spectrum, and clear differentiation trajectory, the hematopoietic hierarchy roadmap was reformed (Figure 4A). The shapes of the three transitional branches were controlled by the expression of key regulatory genes, which functioned similarly to a gradual activation switch. During initial progenitor-differentiation, GATA2, HOPX, and CSF3R regulated the formation of the first branches of the three transitional lineages (Figure 4A, B). Downregulation of *CSF3R* and upregulation of *GATA2* implied that the progenitors were primed to generate oligo-potent progenitor GAPs (Figure 3B). Inversely, the downregulation of *CSF3R* and *GATA2* would prime the MPC committed to CLPs with the upregulation of HOPX and HOXA9. Hence, the separation between NMPs, CLPs, and megakaryocyte/erythroid lineages was determined by GATA2, HOPX, and CSF3R, not GATA1 [4, 26]. That is, *GATA2, HOPX,* and *CSF3R* expression marked the relative boundary between GAPs, CLPs, and NMPs (Figure 4B). The MkE lineages differentiated independently from the other two branches. The expression of *GATA1* and *KLF1* was gradually increased at later mature stages, at which point cells were determined and defined as MEPs (Figure 2B). CSF3R was upregulated in the original MPCs and NMPs but was nearly absent during the later stages of MEPs (C1) and CLPs (C7) (Figure 2B and 4B). GATA2, HOPX, and HOXA9 expression was restrained when the progenitors were primed to generate the NMP branch (Figure 3B, C). Cytokine receptors were co-expressed or co-silenced with relevant key lineage regulators, indicating that multiple genes regulated the progenitor lineage-priming process. During the differentiation of the MkE lineages, the expression levels of unique genes, such as *GATA2*, *CPA3*, *CNRIP1*, and *SLC40A1*, gradually increased. We also identified genes with the opposite expression pattern that were specific among MkE lineages (C1, C6, and C11) and other progenitors, such as *SPINK2*, *C1QTNF4*, *NPDC1*, *S100A10*, and *SELL*. Enhanced MEP differentiation resulted in the formation of the second branch of mature MEPs. Toward the final maturation stages of MEPs, KIT, GATA2, KLF1, or LMO4 expression guided the differentiation of progenitors to megakaryocyte–erythrocyte (Pro-ME) cells or mast cell and basophil progenitors (MBPs) (Figure 4A). The receptor of TPO, MPL, was absent in Pro-ME, indicating that megakaryocyte progenitors were more likely originated from MEPs (C1) but not co-segregated with erythrocytes in the Pro-ME cluster. Additionally, *LMO4*, *MS4A2*, and *TPSB2* were uniquely upregulated in MBPs. The hemoglobin subunit delta was identified as a protein involved in early erythrocytic development. Mini-chromosome maintenance protein genes, encoding components of the MCM2–7 complex, were highly expressed in Pro-ME cells and NMPs (Table 1). The MCM complex is the replicative helicase that plays an essential role in the eukaryotic DNA replication-licensing system [27].Our results indicate that the status of the two lineage progenitors (NMPs and Pro-MEs) may switch from differentiation (commitment) to proliferation. Finally, the continuous differentiation process of neutrophils was directly primed from MPCs and underwent two commitment stages before transfer to proliferation (Figure 4A).

**Figure 4.**
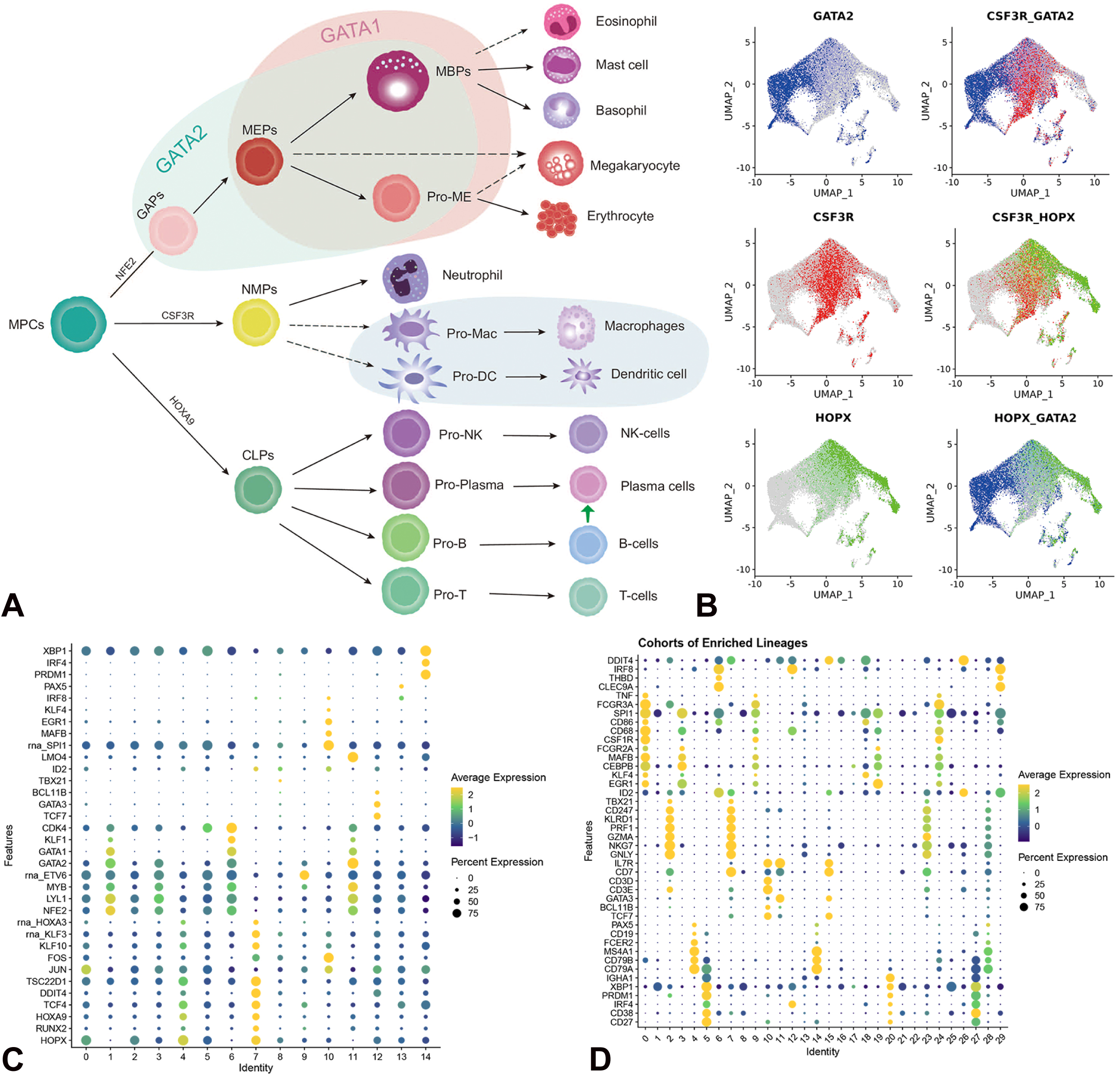
Reformed hematopoietic hierarchy of progenitor cell differentiation. A, Reformation of the **hematopoietic hierarchy** based on an scRNA transcriptional atlas. In the new hematopoietic hierarchy, the MPCs are simultaneously primed into three transitional progenitor cell types, which subsequently mature into diverse lineages. The first progenitor cell type is the CLPs, which differentiate stepwise into T, B, and NK progenitor cells. The second progenitor cell type, GAPs, initially undergoes continuous differentiation into intermediate-stage MEPs through processes controlled by GATA1 and GATA2. During the later stages, MEPs differentiate into two directions, controlled by KIT, KLF1, and LMO4. The high-expression branch differentiates into MBPs. The other branch differentiates into Pro-MEs. The third type of progenitor cells are NMPs, which differentiate into neutrophils and monocytes. The dashed lines represent partial lineage connections that lack strong direct evidence of differentiation relationships. The shaded area indicates the GATA2 and GATA1 expression domains. B, Feature plots illustrating the expression patterns of GATA2, HOPX, and CSF3R in co-segregated progenitors. C, Dot plot illustrating the expression of cell-fate decision-associated TF genes in enriched CD34 progenitors. D, Dot plot illustrating the expression of mature cell marker genes and cell-fate decision-associated TF genes in enriched lineage cohorts. CLP, common lymphoid progenitors; GAP, gene-controlled progenitors; MBP, mast cell, basophil, or eosinophil progenitors; MEP, megakaryocytic–erythroid progenitors; MPC, multipotent progenitor cells; NMP, neutrophilic and monocytic progenitors; Pro-ME, megakaryocytic and erythroid progenitors; scRNA, single-cell RNA; UMAP, uniform manifold approximation and projection.

Our results revealed that the original progenitors of basophils, eosinophils, and neutrophils differed (Figure 4A), consistent with the results of a previous study, which reported that the origin of neutrophils is distinct from that of basophils and eosinophils during lineage commitment [4]. However, the identity of the original eosinophil progenitor was not confirmed. The original progenitor cells were primed and differentiated into various progenitors through gradual activation/deactivation followed by TF accumulation, synergism with key receptor genes, and signal activation. MEP-representative genes gradually accumulated during differentiation, revealing their distinct continuous differentiation characteristics (Figure 2B and 3C). These findings suggest that the hierarchical organization of MkE lineages during hematopoiesis is a continuous process controlled by the activation and suppression of cell-fate decision-associated genes.

During priming of the lymphoid progenitor branch, critical TFs (including *HOXA9* and *HOPX*) were upregulated (Figure 3C, D). Similarly, marker genes of lymphocytic lineages (such as *MS4A1*, *GNLY*, and *GATA3*) were uniquely expressed during specific stages and upregulated stepwise. Certain genes were prominently and specifically co-expressed in lineage-restricted progenitors and CLPs: CLPs expressed *IGHM*, *FCMR*, *BLNK*, and *LTB*; B progenitors expressed *CD79A* and *JCHAIN*; monocyte progenitors expressed *SPI1*, *FOS*, *PECAM1*, and *LST1*; and natural killer (NK) progenitors expressed *NKG7* and *GZMA*. The expression profiles of these genes also indicated the underlying mechanisms of lymphoid lineage commitment. The lymphoid lineage-commitment process is likely a discrete stepwise process, with distinct lineage-committed populations formed during hematopoiesis. This was also observed for monocyte-lineage differentiation into macrophages and dendritic cell (DC) progenitors. We also identified the co-expression of *ETV6, LYL1,* and *NFE2* in NMPs and monocyte lineage-restricted progenitors, resembling the co-expression of *SPI1, FOS, PECAM1*, and *LST1* in monocytes and CLPs (Figure 2C and 3B). However, we were still unable to confirm the original monocyte lineage during hematopoiesis. The commitment of each progenitor lymphoid lineage was controlled by the expression of key regulatory genes, which collectively functioned as a stepwise activation switch. By switching the sequential expression of TFs, which generally showed a synergistic relationship with key receptor gene expression and signal activation, the original progenitor cells were primed and assigned to the subsequent stages of lymphoid-lineage progenitors (Figure 4). This study is the first to clear define the initial primed transitional progenitor and reformed the hierarchy, effectively reconciled the puzzle concerning pathways and models of hematopoiesis commitment.

### Identification of cell fate decision-associated genes in HPC maintenance and lineage commitment

Cell-fate decision-associated factors were identified in different lineages, including *NFE2, LYL1, MYB, CDK4, ETV6, TESPA1, ZBTB16*, and *GATA2* in myeloid progenitors and *RUNX2, HOXA9, JUN/FOS, TCF4, DDIT4, HOPX, KLF10, HOXA3*, and *TSC22D1* in lymphoid progenitors (Figure 4C). Notably, these key factors were determined to play contrasting functional roles during HPC differentiation. HPCs differentiated into diverse myeloid progenitors by enhancing the expression and functions of *NFE2, LYL1, MYB, CDK4, ETV6, TESPA1*, and GATA2 while suppressing the expression and functions of *RUNX2, HOXA9, TCF4, DDIT4, HOPX, KLF10, HOXA3*, and *TSC22D1*. Together with the activation of other receptors and unique TFs (such as CSF3R in neutrophils and CSF2RB in MkEs) in each progenitor lineage, the entire hematopoietic lineage-commitment process was precisely controlled and occurred gradually (Table 1 and Figure 4A, C).

To identify the essential genes involved in maintaining HPC states, we compared the expression levels in three cohorts of bone marrow cells, control PBMCs, and CD34+ single cell subsets. We classified the progenitor-maintaining genes using the following criteria: first, the gene must be pan-expressed across most PBMC and bone marrow HPC clusters but should be excluded once it serves as a specific lineage fate decision gene (such as GATA1 for MkE lineages); second, the expression of this gene should be limited to progenitor cell types. Such genes, pan-expressed in both progenitor and mature cells, would be considered HPC functionally-related genes but would not be associated with progenitor maintenance. In this regard, our data revealed that *RUNX1* regulated the differentiation of most progenitors (Figure 3C), whereas *RUNX2* primarily regulated the differentiation of lymphoid progenitors. We also found that ITGA4 (CD49d) was highly expressed in most progenitors, which is important for molecular regulation during HSC homing [28]. Functional HSC-marker genes, including *PTPRC*, *HMGB1*, *FOXP1, CD74*, *CD37*, *PFN1*, *TXN*, *ZFP36L2*, *SERPINB1*, *TMSB4X*, and *HSP90AA1,* were broadly expressed in most progenitor cells but were not specific (Table 1 and Figure 1F). These genes may mediate the self-renewal and mobilization, but not differentiation and maintenance, of HPCs. Additionally, two new genes (*PPIA* and *EIF1*) were identified, which were similar to previously reported genes as they exhibit pan-expression in progenitors and were highly expressed in mature subgroups of peripheral blood cells.

*XBP1, SOX4, CD82, SPINT2, IMPDH2, TSC22D1, SPI1* [14], and *ETV6* were also widely expressed in most progenitors and progenitor-differentiated cells; however, their expression was relatively absent in all other cell types and control cohorts (Figure 1F and Table 1). Moreover, *ERG, BCL11A, LMO2*, and *RUNX1* were widely expressed in approximately 50% of the cells in each progenitor subgroup. These genes, similar to EGFL7 and CYTL1 (Figure 1F), were more highly or specifically expressed in the bone marrow and progenitor cells than in mature PBMCs (Figure S5), which highlights their significance in regulating the HPC state for maintenance and differentiation.

### Identification and validation of fate decision-associated genes in each redefined subgroup

To identify cell-fate decision-associated genes, particularly those encoding TFs, advanced analysis was performed on enriched CD34+ subsets. The unique cell-fate decision-associated TFs were identified in each cluster of lineages (Figure 4C). Four key TFs (GATA3, TCF7, BCL11B, and DDIT4) were identified in T progenitor cells (Figure 4C). These four TFs were critical for cell fate decisions during T-cell lineage differentiation (Table 1 and Figure 4C). Previous findings suggest that *TCF7* and *GATA3* are critical TFs for T-cell development and differentiation [29, 30], which is highly consistent with our redefined characterization of T progenitor cells. We observed that LTB was widely expressed by CLPs, Pro-T, and Pro-B progenitor cells and that it was upregulated during the later stages of lymphoid progenitor differentiation. Two distinct types of B progenitor cells were identified and redefined as expressing different cell-fate decision-associated TFs and marker genes in C13 and C14. Two TFs (*PRDM1* and *IRF4*) were specifically expressed in C14, whereas *PAX5*, *IRF8*, and *BACH2* were specifically expressed in C13 (Figure 4C). *XBP1* was widely expressed in both clusters and upregulated in C14. C14 comprised the progenitors of plasma cells, whereas C13 represented classical B progenitor cells (Table 1, Figure 4C). These data indicate that the development and differentiation of B cells occur during the early progenitor stages. Plasma cells are thought to represent the final stage of B cell proliferation and are incapable of further proliferation [31]. However, our scRNA data confirmed the existence of plasma progenitor cells in whole blood samples. The immunoglobulin variable (IGV) genes were expressed primarily by the progenitors of C14 cells, with rearrangements in the *IGHV*, *IGKV*, and *IGLV* genes having already occurred during the progenitor state of plasma cells. These six TFs, *PRDM1*, *IRF4 PAX5*, *IRF8*, *BACH2,* and *XBP1,* play critical roles in the cell fate decisions of B lineage cells. HOPX was significantly highly expressed in progenitor and mature NK cells but was not expressed in MkE lineages, suggesting that HOPX was a crucial TF for NK cell development (Figure 4C).

Monocyte progenitor cells in C10 were identified and redefined. One of the most highly expressed genes in monocyte lineages was *IFI30* (Figure 2F), which is related to the interferon-gamma signaling pathway [32]. *SPI1*, *PECAM1*, and *LST1* were widely expressed in most monocyte progenitors, suggesting that *SPI1* and *LST1* are critical genes for monocyte progenitors. The monocyte progenitors were controlled by various cell-fate decision-associated TFs, including *KLF4, EGR1, MAFB, CEBPB, BCL6, IRF8*, and *DDIT4* (Table 1 and Figure 4C). To identify and validate the fate decision genes among the two monocyte cell types (macrophages and DCs), TF expression patterns were reanalyzed in the control cohorts. The macrophage cells expressed high levels of *EGR1*, *CEBPB*, and *MAFB* but did not express *IRF8* or *DDIT4* (Figure 4D and S5). Furthermore, the macrophages included FCGR3A^+^ cells and showed high levels of CSF1R and TNF expression. The DC cells positively expressed *CLEC9A, ANPEP*, and *THBD* (Table 1) and highly expressed *IRF8, BLC6*, and *DDIT4* (Figure S5). The fate decision of macrophages and DCs were determined by these TFs.

Validation was conducted in three cohorts of bone marrow cells, control PBMCs, and enriched lineages (Figure 4D and S5). We analyzed the expression characteristics of marker genes and identified TFs of progenitors in control cohorts. MAFB was uniquely expressed in mature macrophages, TBX21 was uniquely expressed in mature NK cells, and PRDM1 was uniquely expressed in mature plasma cells (Figure S5). These findings were highly consistent with the fate-decision identification in redefined progenitor clusters. Results showed that the cell-fate decision-associated TFs were uniquely expressed in their respective differentiated mature cell types. Furthermore, these cell-fate decision-associated TFs showed more accuracy and efficiency in mature immune cell classification (Figure S5). Overall, the characteristics of cell-fate decision-associated TFs in lymphoid, monocyte, and MEP redefined progenitor lineages were highly consistent with the mature cell types. Cell-fate decision-associated TFs in each branch of the hematopoietic hierarchy were clearly recovered.

### Signaling pathways associated with commitment in each lineage

Diverse pathways affect and regulate HPC differentiation. Our analysis showed that most progenitors expressed high levels of *JUN/FOS* (*AP-1*), *JUNB*, and *JUND* (Figure 2C and 3F). These data indicate that the JUN-related signaling pathway is critical for the differentiation, mobilization, and maintenance of HPCs. MkE clusters expressed high levels of *DEPTOR* (Figure S6), which negatively regulates the mTORC1 and mTORC2 signaling pathways [33], indicating that the mTOR signaling pathway should be suppressed during MkE maturation and differentiation. *NFKBIA* was widely expressed in >80% of the progenitor cells, whereas *NFKBIZ* was highly expressed in the progenitors of monocytes and macrophages (Figure S6). *NFKBIZ* is a member of the NF-κB inhibitor family [34], suggesting that the NF-κB signaling pathway may be suppressed during monocyte and macrophage maturation and differentiation.

Gene set enrichment analysis of clusters using gene set variation analysis revealed the pathways and signals that affect the differentiation of each branch and progenitor subgroup (Figure 5A). The transforming growth factor-beta (TGF-β) signaling pathway was enriched in the original progenitor cells (C0, MPCs), demonstrating its importance in maintaining the quiescent state of HPCs [35]. Moreover, the Notch, interferon alpha and gamma response, TGF-β, and TNF-α signaling pathways were enriched, indicating their potential importance in maintaining monocyte progenitors (Figure 5A). However, expression of the TNF-signaling gene set was suppressed in MEP progenitors (Figure 5A). A previous study has revealed the novel role of Notch in regulating monocyte survival during hematopoiesis [36]. Heme-metabolism and hedgehog-pathway gene sets were enriched in erythroid progenitors (GAPs). Activation of TGF-β and the hedgehog pathway gradually decreased during the committed stage of erythroid progenitors (MEPs), whereas oxidative phosphorylation increased. The NOTCH signaling pathway was highly enriched in MBPs. Hallmark cell proliferation-related gene sets related to the G2M checkpoint, E2F targets, MYC targets (V1 and V2), and the DNA repair pathway (defined by the molecular-signature database) were highly enriched among erythroid and neutrophilic progenitors (Figure 5A). The presence of cell proliferation signaling in erythroid (C6) and neutrophil (C5) progenitors indicates that these cells completed differentiation and tended to proliferate. The IL6/JAK/ STAT3 signaling pathway was enriched in CLPs. The Wnt/β-catenin signaling pathway was enriched in T cell progenitors (C12), which was previously reported to regulate peripheral T cell activation and migration [37]. Moreover, interferon-response gene sets were enriched in B cells and monocyte progenitors.

**Figure 5.**
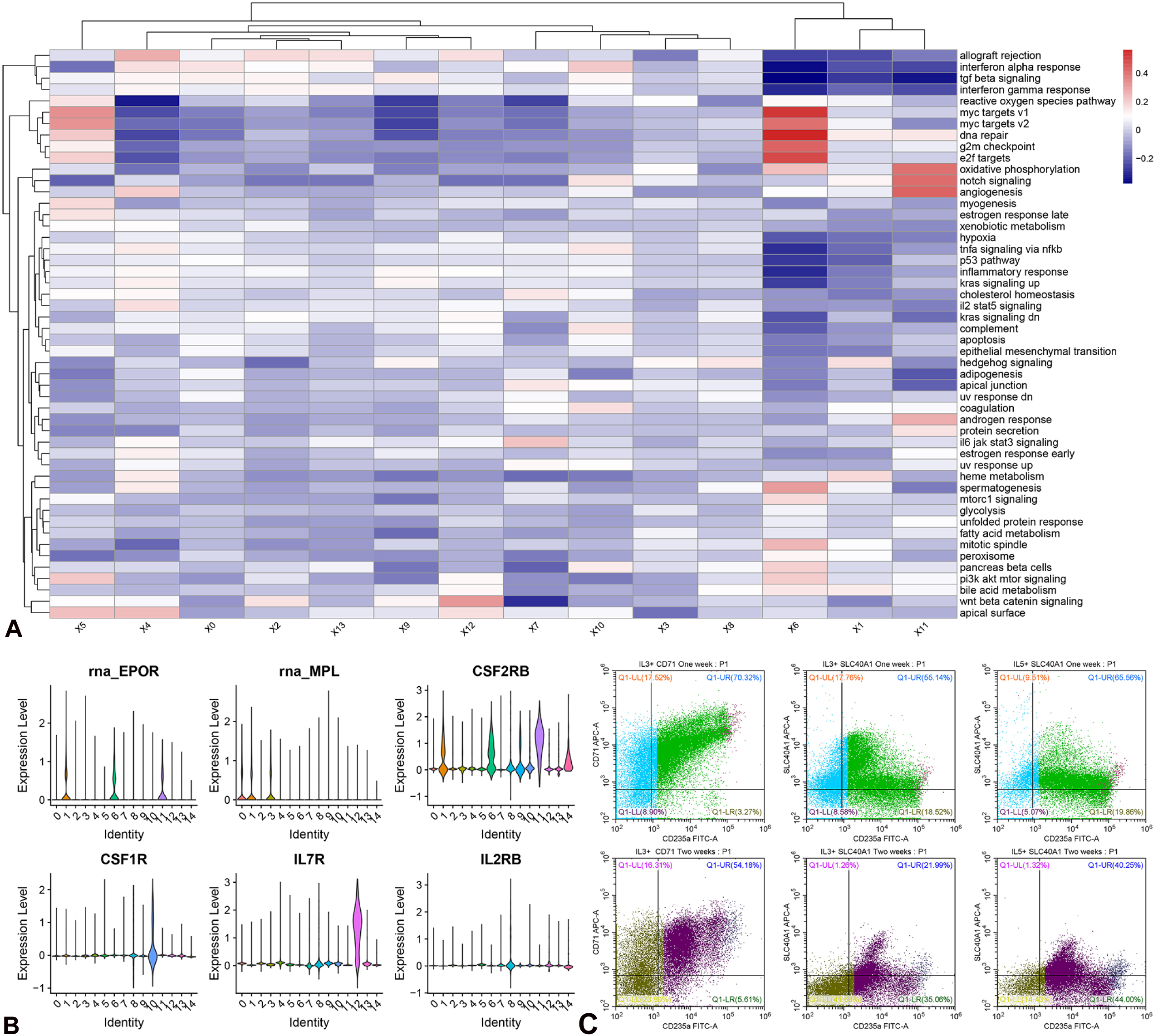
Gene set enrichment analysis and flow cytometry validation of the redefined hematopoietic progenitor cells. A, Heatmap plot showing gene set enrichment analysis results for GSVA clusters. B, Violin plot illustrating the expression of receptor genes. The receptors encoded by *CSF1R*, *CSF2RB*, and *MPL* were specifically expressed in the redefined progenitor subgroups. C, D, Flow cytometric analysis demonstrating the frequency of CD235A^+^ cells and the distribution of SLC40A1^+^ cells during 1 (C) or 2 weeks (D) of primary culture. Higher percentages of CD235A^+^ cells were detected in the IL-5 groups than in the IL3 groups. The plots illustrate the dynamic change in the distribution of SLC40A1^+^ cells, which was more efficient than that of CD71 during distinct erythrocyte cell stages in *in vitro* culture. GAPs, gene-controlled progenitors; GSVA, gene set variation analysis; HPCs, hematopoietic progenitor cells; MEPs, megakaryocytic–erythroid progenitors.

### Functional validation and primary culture confirmation of the reliability of the redefinition of HPCs

The functions of HSC-fate decision-associated genes have been validated previously [14, 38]. We hypothesized that the phenotypic results of functional studies would be consistent with or explained by our HPC definitions. Furthermore, the reformed model of the hierarchical organization of hematopoiesis can be used to interpret conflicting or unclear results. For example, KLF1 and GATA1 are key cell-fate decision-associated TFs involved in erythroid and megakaryocytic differentiation [26, 39]. Functional studies have confirmed that GATA1 progenitors generate mast cells, basophils, megakaryocytes, and erythroid cells [26, 40], while their absence results in the production of monocytes, neutrophils, and lymphocytes [26]. Our transcriptional atlas revealed that GATA1 was uniquely expressed in the MEP but not in monocyte, neutrophil, or lymphocyte progenitors (Figure 2B). These expression characteristics explain the functional study results, indicating that the absence of GATA1 halts MkE lineage differentiation. Conversely, the absence of GATA1 causes progenitor cells to differentiate into NMPs and monocytes. In primary cultures, HPCs differentiated into erythroid and mast cells under the same *in vitro* conditions, utilizing IL3 and SCF; this indicates that both cell types originated from the same lineage. These results confirmed the reliability of the reformed hematopoietic hierarchy of the MkE lineage.

Macrophage colony-stimulating factor (M-CSF) is critical for macrophage survival during *in vitro* primary cultures [41]. This observation supports our transcriptional atlas findings, in which the M-CSF receptor, CSF1R, was only expressed in monocyte progenitors and mature macrophages (Figure 5B). Stem cell growth factor (SCGF), also known as CLEC11A, is a secreted glycoprotein that exhibits erythroid burst-promoting activity in the absence of the SCF/c-Kit ligand [42, 43]. Our results showed that SCGF was highly expressed in NMPs and MEPs, whereas CSF3R, a G-CSF receptor, was highly expressed in NMPs (Figure S6). SCGF cooperates with GM-CSF or G-CSF and enhances the growth of granulocyte-macrophage colonies; however, if only SCGF is present in the culture, erythroid-burst activity is promoted [42, 43]. The expression characteristics of CSF3R and G-CSF strongly suggest that SCGF, together with G-CSF, activates and induces the differentiation of neutrophil and macrophage progenitors, suppressing MkE lineage generation. The expression of the IL15 receptor IL2RB in NK progenitors was unique (Figure 5B). The common beta chain receptor CSF2RB of IL3, IL5, and GM-CSF was uniquely expressed in erythroid lineage progenitors. Results based on the primary HPC cultures showed that progenitors could differentiate into a higher percentage of CD235A^+^ cells after treatment with IL-5, erythropoietin (EPO), and SCF than with IL3 (Figure 5C). The expression patterns of IL3RA and CSF2RB in our atlas revealed that IL3 can non-specifically bind to both receptors and induce progenitor differentiation into other types of lineages (Figure 5B). Moreover, flow cytometric results showed that the expression of the identified marker *SLC40A1* gradually decreased and that its expression became more unique throughout erythroid differentiation (Figure 5C). These results validated the expression patterns of markers unique to MkE lineages. In addition, IL7R, an essential driver of T-cell development and homeostasis, was solely expressed in T-cell progenitors, whereas *IL2RG* and *IL1B* were expressed across most progenitor subgroups (Figure 5B). IL18 was identified as one of the most essential cytokines in early progenitors (data not shown), playing a critical role in hematopoiesis. *IL1B* transcription was regulated by the TFs *SPI1* and *CEBPB* [44], which were highly expressed in monocyte progenitors (Table 1). Moreover, *ALDH1A1* was highly expressed in erythroid-lineage progenitors (Figure S6), the product of which has previously been shown to participate in retinoic acid synthesis. Retinoic acid is required for progenitor cell maintenance [45]. The expression characterization of these cytokine receptors and genes explains the importance of cytokine ligands in cell survival and proliferation within *in vitro* primary cultures. In summary, the expression characteristics of cell-fate decision-associated TFs and cytokine receptors support that the redefinition of progenitor lineages is verifiable, reliable, and reasonable (Figure 3A).

### Are the definition and hierarchy reliable?

*In vivo* or *in vitro* functional studies can be affected by various conditions, which can’t accurately reflect the true physiological state of original progenitors. And colony forming assay also exhibits obviously defects due to variations in cytokines, samples, and selection methods (see introduction). Most scRNA studies that enrich bone marrow progenitors using flow cytometry enrichment demonstrate inefficient reproducibility, false positives, and subjectivity definitions [2, 12]. Some CD34-positive cells are transcriptionally negative, and subsets are either missing or inconsistently distribution among samples. Fate mapping is nearly impossible for adult original progenitors. Here, we present three aspects of evidence supporting the reliability of our definition and hierarchy.

### Robustly Validating the Definition by a Reliable Atlas

Here, we utilized original progenitors from adult peripheral blood, which faithfully reflect the physiological state of human progenitors. We comprehensively captured the entire differentiation stages of these physiological progenitors. The substantial numbers of cells in each progenitor subset provided a reliable transcriptional atlas (Table S1). By utilizing unsupervised data analysis, we avoided bias and subjective effects on results. The subset distribution reproducibility across sample is exemplary (Figure 2D). Furthermore, the expression patterns of some unique genes in each redefined subset are highly consistent with conventional markers (Table 1). Based on truly physiological original progenitors and a reliable atlas, the scRNA data robustly validates the definition, and is more reliable than definition based on *in vivo* or *in vitro* experiments.

### Straightforward Results Confirm the Reliability of the Hierarchy

Our T-SNE plot clearly and perfectly displayed the three differentiation directions of progenitors after the initial priming step of MPC (Figure 3A). The different stages of the megakaryocytic–erythroid lineages are distinctly distributed in the same directions, expressing key well-known fate decision genes (GATA1,KLF1,TAL1) (Figure 2B).The unique expression characteristics of these well-known genes (GATA1,KLF1,TAL1) confirmed the definition of MEP lineages, and we are the first to identify and define the intermediate stage of GAP(Figure 3A). The subsets of the megakaryocytic–erythroid lineages are adjacent and lack clear boundaries, indicating continuous differentiation trajectories. The gradual and continuous expression characteristics of other well-known genes (GATA2,SLC40A1) also provide robust evidence of the continuous lineage commitment. Furthermore, the unique expression patterns of well-known receptor genes (EPO, MPL, CSF2RB) confirm the definition of MEP lineages, which is consistent with *in vitro* culture. The termination of MEP lineages differentiation yields two type of cells (mast and erythliod), which already validated by numerous studies that mast and erythroid cell are generated under the same culture conditions, and further validation is needn’t.

### Physiological Evidence

The megakaryocytic–erythroid lineage commitment process is continuous, aligning with the constant production of erythroid cells every second in the blood. While the commitment process in the lymphoid lineage is discrete and stepwise, reflecting the obviously increase in numbers and heightened activity of lymphoid cells specifically in response to infections or pathological conditions (adaptive immunity). The continuous differentiation process of neutrophils is the shortest in the hematopoietic hierarchy (only two stages) (Figure 3A, C), which also matching with physiological phenomenon. Neutrophils are the primary and rapid responders during infections of innate immunity and can quickly generate an enormous number of mature neutrophils for immune defense [46]. The shortest differentiation trajectories of neutrophils enable a rapid and robust response against infection. In summary, the reformed hierarchy of hematopoiesis perfectly explains the physiological phenomenon of immunity. Often, the truth is simple, yet we have been confused so long time.

## Discussion

Overall, we redefined the characteristics of the original progenitor lineages and identified specific cell-fate decision-associated TFs and marker genes critical for hematopoiesis. This study is the first to define the initial primed transitional progenitor using reliable TFs and markers, effectively solving the century-old puzzle concerning whether CMPs, GMPs, or LMPPs exist. In addition, we reformed the comprehensive hierarchy of hematopoiesis that reconciled previous models and explained the conflicting results among functional studies [4, 16, 47].

During the first commitment stage of hematopoiesis, the initial hematopoietic progenitors are simultaneously primed into three transitional progenitor branches. Our findings provide detailed insights into HPC lineages, revealing the molecular mechanism underlying their commitment and multipotent differentiation. Our scRNA atlas provides an integrated landscape of the heterogeneity, development, and hierarchy of adult peripheral-blood progenitor cells. Collectively, the results of this study represent significant breakthroughs in the fundamental discipline of hematopoiesis, overcoming disciplinary challenges that have plagued the field for more than half a century.

Currently, no consensus has been reached regarding the definition of all HPC lineages, owing to inconsistent classifications and conflicting functional outcomes [9, 16]. Here, the comprehensive enrichment of the original HPCs in adult peripheral blood was particularly helpful for establishing clear and reliable HPC definitions. We demonstrated that classical surface markers, including CD38, KIT, THY1, CD49f, and FLT3, were non-specific and inefficient for identifying HPC lineages [2, 12]. Hence, the conventional lineage definition of HSCs using these markers requires improvement. Accordingly, we redefined the lineages using reliable TFs and markers, including initial primed transitional progenitors. Certain unique marker genes of the redefined progenitor subgroups were highly consistent with previous characterizations, such as *KLF1*, *HPGDS*, and *MPO* for MEPs, mast cells, and neutrophils, respectively [22, 48, 49]. We redefined most HPC lineages using unsupervised and inartificial clustering for the first time and provided an unambiguous transcription atlas for each type.

Through our transcriptional atlas, we identified the key fate decision-associated factors at each hematopoietic hierarchy level. The activation of key TFs primes progenitor cells, initiating them to differentiate into downstream-specific committed lineages [50]. The expression of TFs during the original and mature stages of progenitors revealed a basic co-segregation between myeloid–lymphocyte and mast–erythroid development. Additionally, during the later stage, MEPs differentiated into two directions owing to changes in the expression of key TFs (*KLF1*, *LMO4*, and *GATA2*). Our findings highlight the importance of TFs and the characterization of genes involved in hematopoietic-lineage fate decisions, providing novel insights into the molecular mechanisms underlying the hematopoietic hierarchy and addressing the inconsistencies among the *in vitro* and *in vivo* studies of progenitor cells [26]. The expression characteristics of known cell-fate decision-associated TFs, such as *NFE2*, *TAL1*, and *LYL1* [51–53], provide complementary evidence that the redefinition of progenitor lineages is verifiable, reliable, and reasonable. Furthermore, we found that changes in the expression of unique TFs were independent and strictly lineage-affiliated, contradicting previous conclusions [54]. Previously, induced pluripotent stem cells were established by activating four TFs [55], and *GATA1* and *TAL1* have been validated in reprogramming megakaryocytes and erythroid cells [56, 57]. To our knowledge, this is the first study to clearly identify the cell-fate decision-associated TFs in each branch of the adult peripheral-blood hematopoietic hierarchy. In turn, precise control of the priming and differentiation of HPCs can be achieved by artificially and precisely controlling these key TFs and receptor signals, which we refer to as the Artificial Operation of HPCs.

### CSF3R, the G-CSF receptor [58], was primarily expressed in the original clusters

(MPCs). However, the molecular mechanisms underlying G-CSF induced-enhancement of hematopoiesis remain unclear. Our study reformed the hematopoiesis hierarchy based on the scRNA analysis of unperturbed and G-CSF-mobilized progenitor lineages in peripheral blood. G-CSF directly stimulated downstream activation signals in HPCs by binding to CSF3R, resulting in the conversion of quiescent-state HPCs into activated progenitors. These HPCs were then primed and committed into various progenitors, after which they further differentiated into mature myeloid and lymphocyte cells. The key TFs and cytokine regulators synergistically controlled hematopoietic-lineage commitment, which was validated using primary culture, lineage formation, and reprogramming studies [14]. However, the mechanisms underlying lineage commitment raise certain questions regarding the following concepts: how key TFs are activated during the lineage-commitment process; whether the TFs are activated independently (internally) or via cytokine-signaling cascades (externally); and which factors regulate the activation of lineage-fate decision-associated factors (TFs and cytokine-signaling cascades) that determine subsequent differentiation after the initial priming of HPC with G-CSF. Generally, environmental factors and epigenetic modifications also play key roles.

The following three aspects strongly support the definition and classification of HPCs as distinctly verifiable, reliable, and reasonable. First, the unique markers were highly consistent with the characteristics of most progenitor subgroups. Second, the redefinition addresses the conflicting results reported in previous functional studies [26]. Third, the expression profiles of distinguished cytokine receptors precisely explain the phenomena observed in primary cultures of progenitor subgroups, such as CSF3R and CSF1R in neutrophil and monocyte cultures, respectively [59].

Recent findings have suggested that HSC lineage commitment is a continuous process [2, 13], contradicting the classical stepwise hierarchy of hematopoiesis [60]. However, owing to HSC heterogeneity and the conflicting definitions of HSC subgroups, the mechanism underlying this hierarchy is enigmatic, resulting in inconsistent and incompatible established hematopoietic hierarchies [8]. Our elaborate and comprehensive transcriptional atlas of HPCs provides accurate definitions and a new biological perspective on hematopoiesis. The clear definition and characteristics of HPCs showed that both stepwise and continuous processes occurred during lineage commitment, which can be distinguished by the expression of unique TFs and marker genes. Bias during HPC differentiation gradually accumulated through the activation of multiple lineage-specific TFs, such as *KLF1* and *GATA2*, in MEP cells. The MkE lineage commitment process is continuous, corresponding with the constant production of erythroid cells in the blood [61]. In contrast, the lymphoid lineage-commitment process is discrete and stepwise, reflecting the obvious increase in numbers and heightened activity of lymphoid cells in response to infections or pathological conditions (adaptive immunity). Neutrophils have the shortest differentiation trajectories, which enable a rapid and robust response against infection. These findings facilitate mapping the entire set of transcriptional signaling and pathways involved in lineage commitment and provide strong evidence of a heterogeneity gradient among lymphoid and MkE lineage progenitors.

Previous studies have proposed updated models of the human hematopoietic hierarchy [4, 15]. The major difference between these models and our hierarchy is that we clearly defined the first priming (three directions) and transitional stages (GAP) of the commitment process. We confirmed that initial hematopoietic progenitors were simultaneously primed into three transitional branches of megakaryocyte/erythroid, lymphoid, and neutrophilic progenitors during the first differentiation stage of hematopoiesis. LMPPs, GMPs, and CMPs likely represented a heterogeneous mixture of transitional progenitors. CSF3R was positively expressed in the early stages of erythroids (GAPs) but not in the lymphoid lineages (CLPs). The reformed hematopoietic hierarchy further explains how diversified progenitor cells are primed, induced, and differentiate into all lineages of immune-system cells. The HPC transcriptional atlas provides a new perspective and an adequate explanation of how the immune system is reconstituted in adults. This reliable redefinition and hierarchy will clarify clonal hematopoiesis dynamics. Controlling the initial stage and generation of cytokines, particularly by suppressing the initial cytokine-mediated activation of cascade processes, could inhibit or reduce the effect of subsequent cascade activation. This approach may be a novel therapeutic strategy to control the excessive activation of cytokines and prevent cytokine storms [62, 63]. Thus, our reformed taxonomy and hierarchy will improve immune monitoring in healthy and diseased states and precise prevention and treatment.

This study has certain limitations. First, granulocyte, neutrophil, and monocyte progenitors were identified as originating from the same myeloid progenitors [9]. However, our results showed that the cell-fate decision-associated TFs and characteristic genes significantly differed between these cells, implying that the original association between these lineages should be reconsidered. Our results further showed that monocyte lineage-restricted progenitors and CLPs had similar expression patterns. We speculate that monocyte progenitors could also originate from CLPs or have an independent origin. Although we identified cell-fate decision-associated TFs in two types of B progenitor cells, further investigations are required to determine whether both types of progenitor cells were derived from the same or independent original cells. Second, our enrichment method might have caused the loss of some intermediate-state progenitors, such as the committed megakaryocyte progenitor. Given that our entire atlas was established based on unsupervised dimensionality reduction and clustering from a statistical approach, its accuracy and efficiency remain imperfect. Artificial intelligence and deep learning could provide an accurate, reliable, and distinguishable definition for each state and all progenitor subgroups during hematopoiesis. Moreover, DNA barcoding and the clustered regularly interspaced short palindromic repeats (CRISPR)/CRISPR-associated protein 9 (Cas9)-mediated genome-editing system could help evaluate the clonal dynamics of progenitor cells during hematopoiesis [64].

In conclusion, our results enabled us to reform the comprehensive hierarchy of hematopoiesis and identify key regulators that control hematopoiesis in human adult peripheral blood. Our new model accurately defines a hierarchy that provides biological insights into the hematopoiesis and lineage commitment of HPCs, aiding future research in cell engineering, regenerative medicine, and disease treatment. The reformed lineage-commitment results provide critical insights into the etiology and treatment of hematologic disorders. Moreover, this hematopoietic hierarchy has the potential to achieve artificial, precise, and controlled progenitor cell differentiation and targeted treatment of immune diseases.

## Acknowledgments

We acknowledge Professor Chen Qiu, Rongchang Chen, and Yingyun Fu for supporting this research. Moreover, we are grateful to the participants for their involvement in this study. We would like to express our appreciation for the laboratory support provided by the Key Laboratory of Shenzhen Respiratory Diseases (ZDSYS201504301616234). We also thank Chi-Wai Man of the Chinese University of Hong Kong for manuscript revision. The study was supported by the National Natural Science Foundation of China (grant number 21807072).

## Authors’ contributions

Y.Y. conceived the original idea, designed the study, and performed HPC-enrichment experiments. Y.Y. and L.C. conducted the bioinformatics and statistical analyses of the sequencing data. Y.Y. and Q.S. wrote and revised the manuscript. Q.H., Y.F., and G.L. conducted the sample collection, and S.C generated the schematic diagram. Y.Y. and S.C. conducted the primary culture and laboratory work. All authors approved the final draft of the manuscript.

## Declaration of Generative AI and AI-assisted technologies in the writing process

Not applicable

## Methods

### Sample collection and HPC enrichment

We obtained original peripheral blood HPCs from quiescent donors who had not received any treatment (11 individuals) and mobilized (G-CSF) donors (three individuals). PBMCs and total white blood cells were obtained from 8 mL of peripheral blood according to standard protocols via density-gradient centrifugation with Ficoll® (Cytiva, GE Healthcare). Since lineage commitment is a continuous process, samples collected at any time point can be used to capture all progenitor stages in peripheral blood. Prior to blood collection, mobilized donors received 300 µg of subcutaneous recombinant human G-CSF at 12 h intervals for 5 days. Subsequently, 10 mL of peripheral blood was collected in heparinized tubes from each stem cell donor. Lineage (Lin−, CD3−CD19−CD56−) negative cells were removed using the RosetteSep Enrichment Kit (STEMCELL Technologies, 15272) to enrich the progenitors. The self-designed negative deletion method was applied to enrich the HPCs, enabling the successful capture of the entire spectrum of original HPCs with a sufficient number of cells. Ethical approval was obtained from the ethics committee of the Shenzhen People’s Hospital in adherence with the ethical standards recommended in the 1964 Declaration of Helsinki. Informed consent was obtained from each study participant.

### ScRNA preparation and sequencing

RNA sequencing was performed with human-enriched HPCs (Lin−) purified from 14 donors, including progenitor cells mobilized with G-CSF. Each single-cell suspension was loaded, captured using chromium microfluidic chips, and barcoded with a 10× Chromium Controller (10x Genomics). The barcoded scRNA was reverse-transcribed, and sequencing libraries were constructed using reagents from the Chromium Next GEM Single Cell 5′ Reagent Kit v2, according to the manufacturer’s instructions (10x Genomics). Paired-end sequencing (150 base pairs) was performed using an Illumina NovaSeq 6000 System.

### ScRNA sequencing data processing

The raw sequencing reads of all samples were demultiplexed and aligned to the reference genome hg38 via the Cellranger pipeline. All downstream single-cell unsupervised dimensionality-reduction and clustering functions were performed using Seurat [65]. Progenitor definition and annotation were conducted based on our custom standards without the use of software. GO enrichment analysis of marker genes was performed using the cluster profiler package in R software. The lineage priming trajectories were directly confirmed by progenitor characterization without the need for in-silico prediction. Complementary trajectory analysis was performed using Monocle 2 and Monocle 3 [66]. TF regulation inference and gene regulatory network analyses were performed using Single-Cell rEgulatory Network Inference and Clustering (SCENIC) [67]. Gene set variation enrichment analysis was performed using a non-parametric and unsupervised method of gene set variation analysis [68].

### Integrated analysis with an online single-cell dataset

The online scRNA datasets GSE120221 [69], GSE117498 [12], GSE145668 [70], and GSE181989 [71] were downloaded from the Genomic Spatial Event (GSE) database. Integrated analysis was performed using Seurat. The bone marrow scRNA datasets of 136,489 cells from GSE120221, GSE117498, GSE145668, GSE137864, GSE193138, GSE116256, and GSE181989 were merged. Similarly, the PBMC scRNA datasets of 33,719 cells from GSE226488, GSE168732, and CNP0001102 were merged.

### Cytokine-based primary culture of HPCs

CD34^+^ mononuclear cells were separated from human cord blood samples using Ficoll® (Cytiva, GE Healthcare) and enriched using RosetteSep Human Hematopoietic Progenitor Cell Enrichment Cocktail (STEMCELL Technologies, 15026). Purified CD34^+^ cells were induced to generate MkE-lineage cells in a cytokine-based primary culture. Purified CD34^+^ cells were cultured in StemSpan SFEM (STEMCELL Technologies, 09605) for 21 days at 37 °C with 5% fetal bovine serum under the following two conditions: (1) 10 ng/mL human SCF (PeproTech), 20 ng/mL EPO (MCE), and 10 ng/mL IL3 (MCE); (2) 10 ng/mL human SCF, 20 ng/mL EPO, and 10 ng/mL IL5 (MCE). Cultures were supplemented with 1× insulin–transferrin–selenium–ethanolamine (ProCell) and penicillin–streptomycin–glutamine (Life Technologies). After 2 weeks, human IL-3 and SCF were withdrawn from the culture medium, and an enucleation culture was performed for each group.

### Flow cytometric analysis of erythrocytes

Cultured cells were counted using a cytometer (Count Star) before and after culture. Flow cytometric analysis of surface markers was conducted weekly. Cells were stained with anti-human CD34-phycoerythrin (1:100), anti-human CD235a-fluorescein isothiocyanate, anti-human SLC40A1–Alexa Fluor 647, and anti-human CD71-adenomatous polyposis coli (BioLegend, 1:100) antibodies, followed by analysis using a flow cytometer (Beckman). The antibodies used are presented in Table S2. The results were analyzed using the CytExpert software.

### Glucose-6-phosphate dehydrogenase (G6PD)-activity assay

G6PD enzyme activity in primary erythrocyte cells was detected using the G6PD Activity Assay Kit (GD9650, Leadman, Beijing, China). Suspensions of differentiated CD34^+^ cells were centrifuged at room temperature (25 °C) and counted using a cytometer. Next, the cells were washed with phosphate-buffered saline, resuspended in 200 μL of distilled water, and incubated for 30 min at room temperature to lyse the differentiated erythrocytes and release G6PD. Thereafter, G6PD activity was detected using a biochemical analyzer (Leadman) according to the manufacturer’s instructions (cobas c 702, Roche).

---

Figure S1. Specific expression of genes by HPCs and marker gene enrichment analysis. A, Expression-feature plots of the progenitor markers CD34, HLF and EGFL7, showing that HLF expression was minimal but EGFL7 expression was highly consistent with that of CD34. B, Expression-feature plots of the progenitor markers CD34 and CRHBP, displaying CRHBP as mainly expressed in the early stages of progenitors. C, Gene Ontology enrichment results for progenitor cells. D, Kyoto Encyclopedia of Genes and Genomes (KEGG) enrichment results of signaling pathways in progenitor clusters. E, Gene set enrichment analysis of differentially expressed genes in hematopoietic cell lineages.

Figure S2. Efficiency evaluation and comparison of canonical and newly identified marker genes across datasets. A, UMAP showing the T-SNE plot for the integrated progenitor cells of the aggregated HPC subsets. B, Violin plot showing gene-expression characterization of newly identified genes in HPCs, and displaying CD34-negative cells clustered in the same subgroup, which indicates that these cells are not progenitor cells. The expression patterns of EGFL7, CYTL1, LAPTM4B, and NPR3 are highly consistent with those of CD34 and are more efficient and sensitive than CRHBP in progenitor cell identification. C, UMAP showing the T-SNE plot of the integrated progenitor cells of bone marrow HPC scRNA cohorts. D, Violin plots displaying gene-expression levels in the bone marrow scRNA datasets. HPC, hematopoietic progenitor cell; scRNA, single-cell RNA sequencing. T-SNE: t-Distributed Stochastic Neighbor Embedding; UMAP, uniform manifold approximation and projection.

Figure S3. Re-clustering of early priming-stage clusters of adult peripheral blood HPCs. A, UMAP showing the distribution of each individual atlas, demonstrating exemplary reproducibility across samples. B, UMAP showing the integrated progenitor cells (C0,C2,C3,C4, and C5 of Figure 2A); a common intermediate progenitor stage could not be distinguished among the myeloid lineages. C, Violin plots for early priming-stage progenitors showing no distinct differentiation characteristics among clusters. CMPs could not be distinguished by FLT3, CD38, and IL3RA expression, indicating that CMPs, GMPs, and LMPP were the heterogeneous mixture of various progenitor types. UMAP, uniform manifold approximation and projection.

Figure S4. Pseudo-time analysis among MPC nearby clusters using Monocle2, excluding far away cell clusters. A, Differentiation trajectory of HPCs constructed via pseudo-time analysis. B, Expression trajectory of GATA2, HOPX, and CSF3R along the in-silico pseudo-time trajectories. C, Location of colored clusters in the reconstructed in-silico pseudo-time trajectory tree. D, Pseudo-time analysis showing the three states of the trajectory. E-F, Expression of MEP (E) and CLP (F) marker genes in clusters and states along pseudotime trajectories.

Figure S5. Comparison of gene expression levels across control datasets. Bone marrow and PBMC scRNA datasets were merged. A, T-SNE plot showing the integrated merged bone marrow scRNA datasets. B, Dot plot illustrating gene expression levels of cell marker genes and cell-fate decision-associated TF genes in bone marrow cohorts. C, T-SNE plot showing the integrated merged control PBMC scRNA datasets. D, Dot plot illustrating cell marker gene and cell-fate decision-associated TF gene expression levels in bone marrow progenitors. PBMC, peripheral blood mononuclear cell; scRNA, single-cell RNA.

Figure S6. Violin plot illustrating the expression levels of genes in enriched CD34 positive clusters.

