## supplemental figures and table for "Redefining hematopoietic progenitor cells and reforming the hierarchy of hematopoiesis"

### Supplementary Figures legends

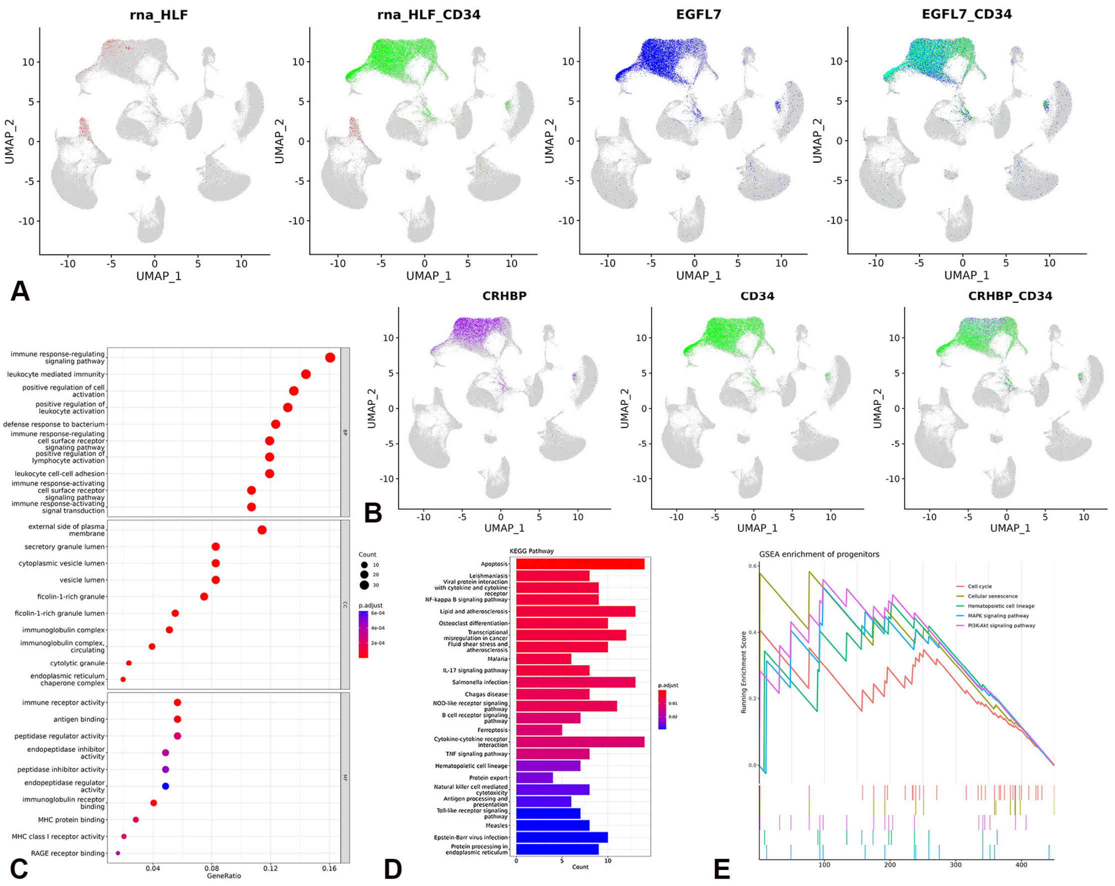


Figure S1. Specific expression of genes by HPCs and marker gene enrichment analysis. A, Expression-feature plots of the progenitor markers CD34, HLF and EGFL7, showing that HLF expression was minimal but EGFL7 expression was highly consistent with that of CD34. B, Expression-feature plots of the progenitor markers CD34 and CRHBP, displaying CRHBP as mainly expressed in the early stages of progenitors. C, Gene Ontology enrichment results for progenitor cells. D, Kyoto Encyclopedia of Genes and Genomes (KEGG) enrichment results of signaling pathways in progenitor clusters. E, Gene set enrichment analysis of differentially expressed genes in hematopoietic cell lineages.

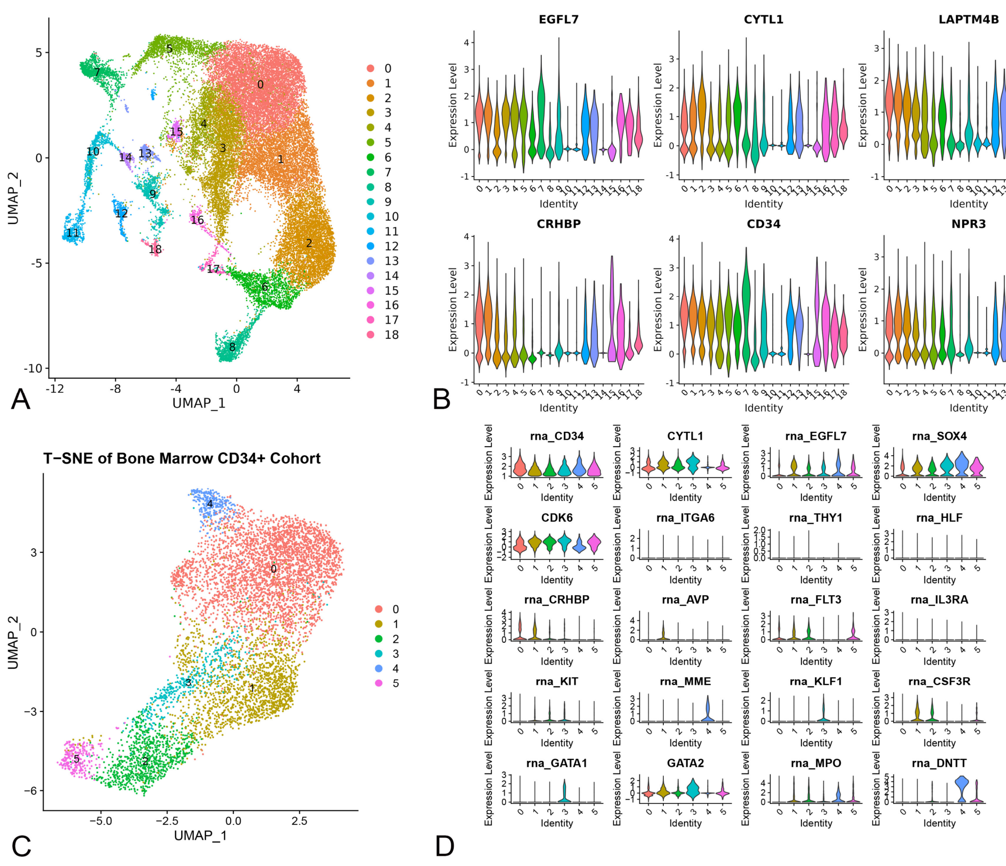


Figure S2. Efficiency evaluation and comparison of canonical and newly identified marker genes across datasets. A, UMAP showing the T-SNE plot for the integrated progenitor cells of the aggregated HPC subsets. B, Violin plot showing gene-expression characterization of newly identified genes in HPCs, and displaying CD34-negative cells clustered in the same subgroup, which indicates that these cells are not progenitor cells. The expression patterns of EGFL7, CYTL1, LAPTM4B, and NPR3 are highly consistent with those of CD34 and are more efficient and sensitive than CRHBP in progenitor cell identification. C, UMAP showing the T-SNE plot of the integrated progenitor cells of bone marrow HPC scRNA cohorts. D, Violin plots displaying gene-expression levels in the bone marrow scRNA datasets. HPC, hematopoietic progenitor cell; scRNA, single-cell RNA sequencing. T-SNE: t-Distributed Stochastic Neighbor Embedding; UMAP, uniform manifold approximation and projection.

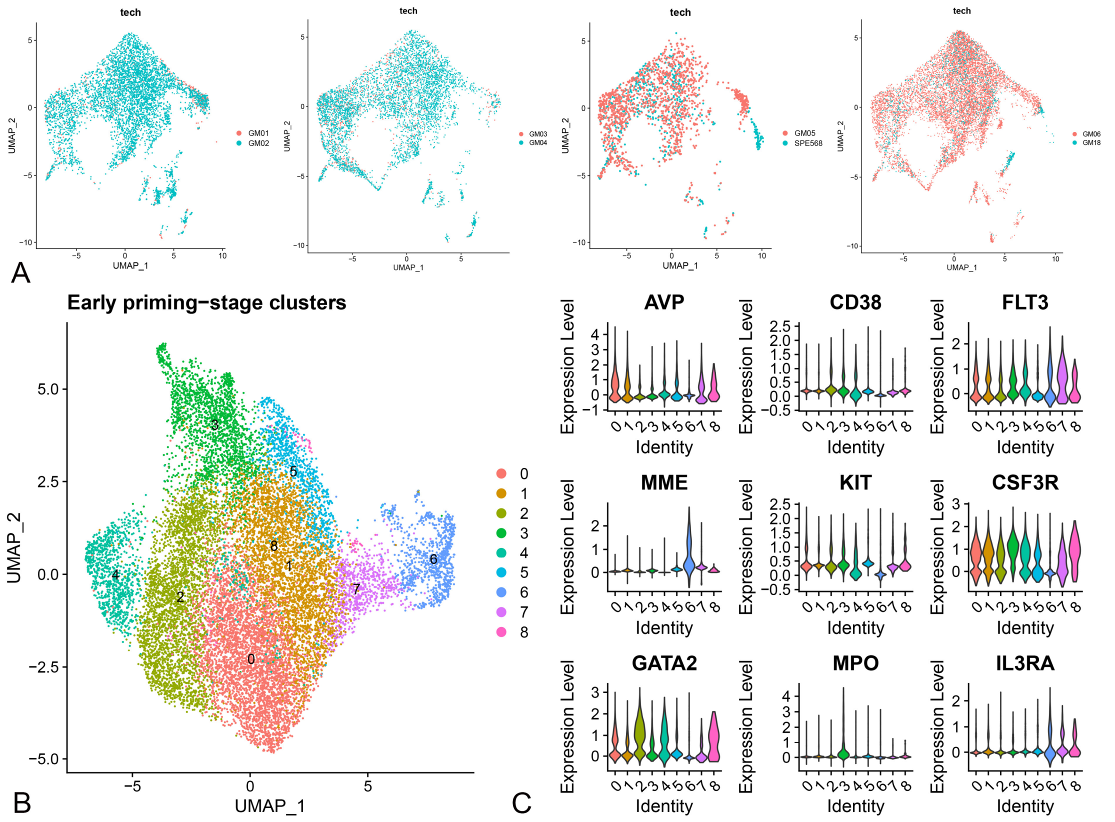


Figure S3. Re-clustering of early priming-stage clusters of adult peripheral blood HPCs. A, UMAP showing the distribution of each individual atlas, demonstrating exemplary reproducibility across samples. B, UMAP showing the integrated progenitor cells (C0,C2,C3,C4, and C5 of Figure 2A); a common intermediate progenitor stage could not be distinguished among the myeloid lineages. C, Violin plots for early priming-stage progenitors showing no distinct differentiation characteristics among clusters. CMPs could not be distinguished by FLT3, CD38, and IL3RA expression, indicating that CMPs, GMPs, and LMPP were the heterogeneous mixture of various progenitor types. UMAP, uniform manifold approximation and projection.

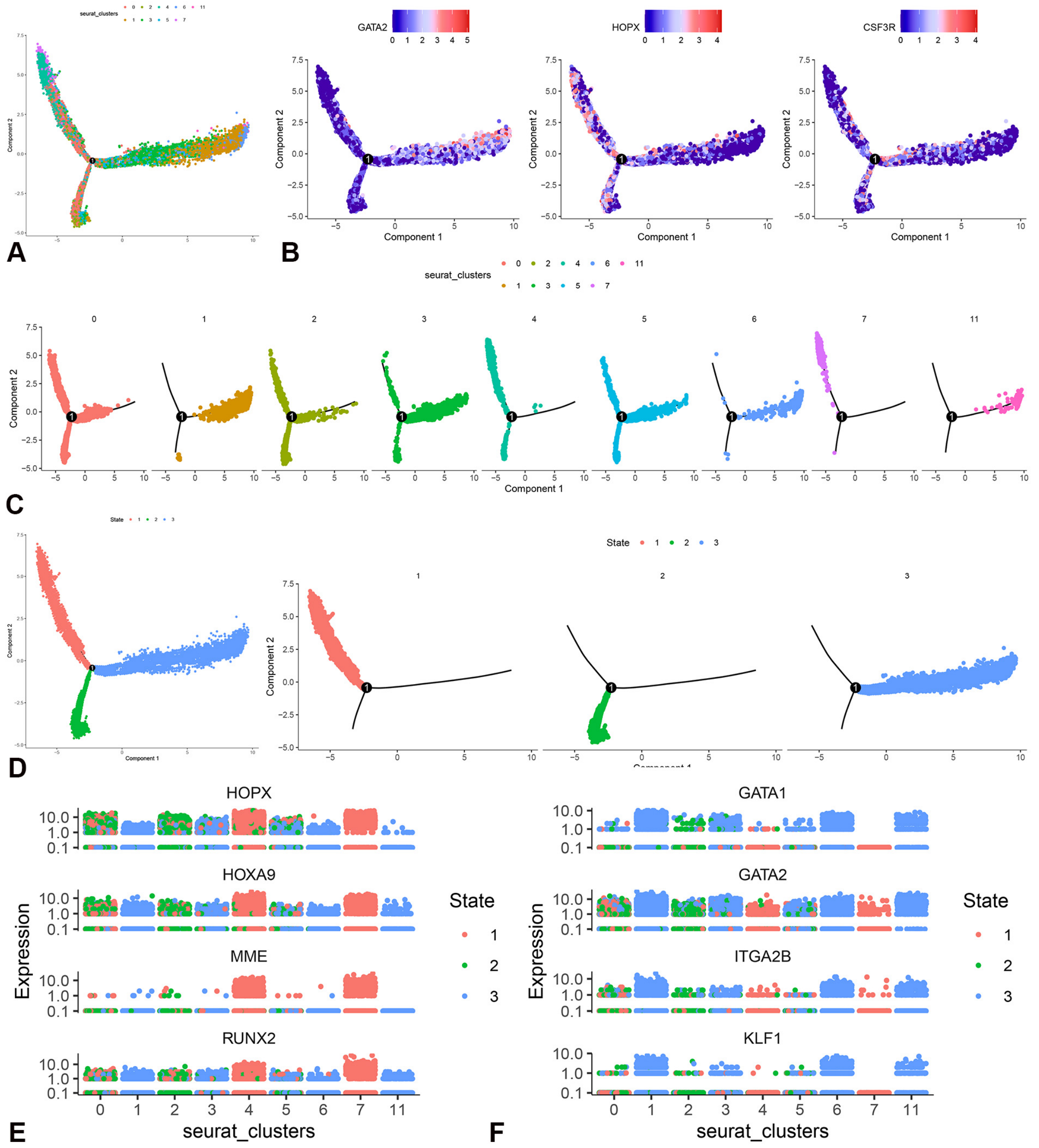


Figure S4. Pseudo-time analysis among MPC nearby clusters using Monocle2, excluding far away cell clusters. A, Differentiation trajectory of HPCs constructed via pseudo-time analysis. B, Expression trajectory of GATA2, HOPX, and CSF3R along the in-silico pseudo-time trajectories. C, Location of colored clusters in the reconstructed in-silico pseudo-time trajectory tree. D, Pseudo-time analysis showing the three states of the trajectory. E-F, Expression of MEP (E) and CLP (F) marker genes in clusters and states along pseudotime trajectories.

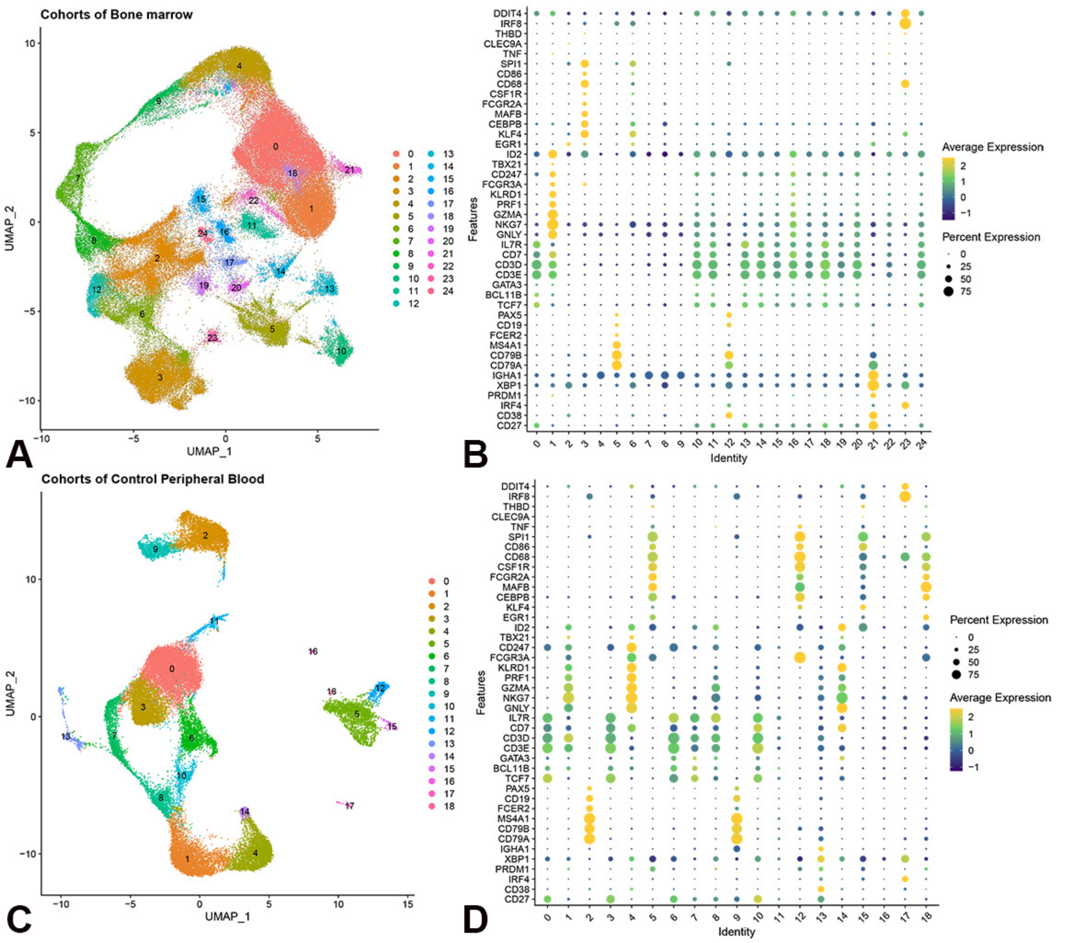


Figure S5. Comparison of gene expression levels across control datasets. Bone marrow and PBMC scRNA datasets were merged. A, T-SNE plot showing the integrated merged bone marrow scRNA datasets. B, Dot plot illustrating gene expression levels of cell marker genes and cell-fate decision-associated TF genes in bone marrow cohorts. C, T-SNE plot showing the integrated merged control PBMC scRNA datasets. D, Dot plot illustrating cell marker gene and cell-fate decision-associated TF gene expression levels in bone marrow progenitors. PBMC, peripheral blood mononuclear cell; scRNA, single-cell RNA.

Table S1. The cell numbers of each cluster among HPCs datasets

| Clusters | C0 | C1 | C2 | C3 | C4 | C5 | C6 | C7 | C8 | C9 | C10 | C11 | C12 | C13 | C14 |
|---|---|---|---|---|---|---|---|---|---|---|---|---|---|---|---|
| Cell numbers | 6445 | 3990 | 3714 | 3276 | 2015 | 1705 | 1177 | 767 | 503 | 452 | 410 | 294 | 294 | 269 | 139 |
